# Elucidating the Sodium Channel-Targeting Anti-Seizure Medication Interactome: A Drug-Target Pathway Analysis

**DOI:** 10.64898/2026.11.13.766542

**Authors:** Srinivasan Ekambaram, Erika T. Axeen, Jaideep Kapur, Howard P Goodkin, Nikolay V. Dokholyan

## Abstract

Epilepsy is a common neurological disorder in both children and adults, primarily managed with anti-seizure medications (ASMs). Sodium channel blockers are among the most commonly used ASMs. These medications exhibit distinct spectra of efficacy and side effect profiles. These likely result from different off-target interactions, protein binding profiles, and subunit selectivity that remain poorly characterized. We developed a neural network-based computational pipeline to systematically elucidate the molecular targets and associated biological pathways of selected sodium channel-modulating ASMs: carbamazepine, lamotrigine, eslicarbazepine, licarbazepine, lacosamide, phenytoin, and cenobamate. Using the DRIFT (Drug–target identification based on chemical similarity) algorithm for proteome-wide target prediction, we identified protein targets for these ASMs. The sodium channel alpha subunit family was the highest-ranked target for every drug except cenobamate, for which it ranked second behind the GABA-A receptor. Pathway mapping in the Reactome database revealed both shared and drug-specific biological processes. Furthermore, molecular docking studies elucidated the atomistic interactions between the drugs and the predicted targets. Specifically, the predicted target landscape varied from eslicarbazepine and licarbazepine (59 predicted targets each) to lacosamide (18). Pathway analysis revealed recurrent enrichment in rapid depolarisation and the interaction between L1 and ankyrins shared across all drugs, alongside predicted off-targets, including the cannabinoid CB1 receptor for phenytoin and the carbonic anhydrases for lacosamide. Molecular docking supported these predictions, with binding scores ranging from -24 to -31 kcal/mol. Overall, this work provides a systems-level view of NaV-targeting ASM pharmacology and a hypothesis-generating framework to prioritize mechanistic and combination studies of these medications in epilepsy.

## Introduction

Epilepsy is a chronic neurological disorder characterized by a propensity for recurrent, unprovoked seizures^1^. Epilepsy can be associated with detrimental effects on the psychological and social aspects of the patient’s life, such as depression, anxiety, and learning difficulties^2^. The International League Against Epilepsy (ILAE) classifies seizures into two biological classes: focal and generalized^3^. Focal seizures originate from a single, localized region of the brain and represent the most prevalent type, accounting for approximately 60% of all epilepsy cases^4^. Conversely, generalized seizures result from aberrant neuronal discharges that rapidly engage networks across cerebral hemispheres. This bilateral activity leads to clinical seizure manifestations such as an abrupt loss of consciousness, falls, or major motor events. Epilepsy is diagnosed after two or more unprovoked seizures or a single unprovoked seizure, where the probability of additional seizures is similar to the general recurrence risk of 60%^5^.

ASMs are initiated following the epilepsy diagnosis, thereby reducing the risk of future seizures^6^. A significant clinical challenge persists in epilepsy treatment, as nearly 30% of patients have drug-resistant epilepsy, even with the availability of newer ASMs^7^. Drug resistance is the failure of two appropriately chosen and tolerated regimens. Although non-ASM options for drug-resistant patients, such as epilepsy surgery, neuromodulation, and ketogenic diets, can be beneficial, a critical void remains in the treatment of epilepsy^8,9^. Moreover, the long-term use of many ASMs is often limited by adverse effects, including sedation, mood and cognitive effects, and bone health^10,11^. Accordingly, there is a continued need to refine ASM utilization through mechanistic optimization and to minimize side-effect profiles.

ASMs can be broadly categorized by mechanisms of action, which target different aspects of neuronal excitability and synaptic transmission^12^. The major classes include drugs that modulate voltage-gated ion channels, enhance inhibitory neurotransmission, or reduce excitatory signaling^13^. Sodium channel blockers, such as phenytoin, carbamazepine, and lamotrigine, stabilize neuronal membranes and prevent repetitive firing by stabilizing the inactivated state of sodium channels^14^. Agents that potentiate GABA-mediated inhibition, such as benzodiazepines and barbiturates, enhance inhibitory tone within neural circuits^15^. Conversely, drugs like perampanel and felbamate act by antagonizing excitatory glutamatergic transmission^16^. Other ASMs, including levetiracetam and brivaracetam, exert their effects by modulating synaptic vesicle protein 2A (SV2A), thereby influencing neurotransmitter release^17^. This mechanistic diversity enables tailored therapy based on seizure type, comorbidities, and individual tolerability, allowing rational polypharmacy when needed by pairing ASMs with varied mechanisms^18^.

Among the major classes of ASMs, voltage-gated sodium ion channel (NaV) blockers are the most widely prescribed medications^14^. Numerous studies have reported on the NaV family, which comprises 9 alpha-subunit isoforms, each encoded by a different gene and exhibiting specific expression in the central and peripheral nervous systems^19^. The gain- and loss-of-function mutations in these genes are linked to various epilepsy syndromes^20^. Although the NaV acting ASMs share common pharmacological properties, their mechanisms of action differ. For instance, carbamazepine binds the fast-inactivated state^21,22^. By contrast, lamotrigine preferentially interacts with inactivated NaVs^23^, whereas eslicarbazepine, the major active metabolite of eslicarbazepine acetate, enhances slow inactivation and shows subtype-dependent activity among NaV channels^24^. These mechanistic differences may contribute to variations in efficacy, tolerability, and, in some cases, worsening of seizures (Figure 1A).

**Figure 1.**
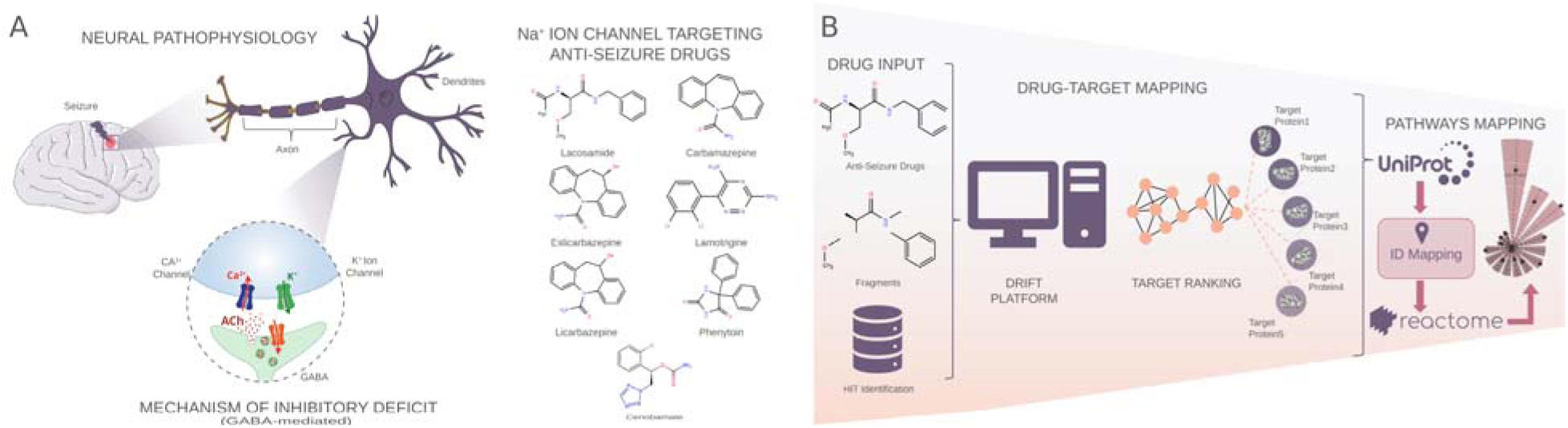
Schematic overview of seizure-related neuronal dysfunction, selected anti-seizure medications, and the DRIFT-based computational workflow. (A) Illustration of epilepsy-related neural pathophysiology, including seizure-focus localization, neuronal architecture, impaired GABA-mediated inhibitory signaling, and dysregulation of calcium and potassium channel function. Structures of seven anti-seizure medications are shown: lacosamide, carbamazepine, eslicarbazepine, licarbazepine, lamotrigine, phenytoin, and cenobamate. (B) DRIFT workflow for target and pathway analysis. Drug structures were fragmented and matched to reference compounds, followed by target prediction using the DRIFT algorithm. Predicted targets were mapped via UniProt identifiers to Reactome pathways for downstream comparative analysis.

Despite their shared classification as NaV blockers, accumulating evidence suggests that these ASMs possess additional, off-target interactions that may contribute to their therapeutic effects and adverse event profiles^25^. In clinical practice, NaV-blocking ASMs are widely used in combination therapies, and clinical evidence indicates that combinations of ASMs with different mechanisms of action (MOAs) are generally more effective than combinations within the same drug class^26^. Moreover, adjunctive therapies such as lacosamide and eslicarbazepine were approved based on pivotal trials in which the majority of patients were already receiving NaV-blocking ASMs, such as carbamazepine or lamotrigine, underscoring their use alongside other sodium channel blockers^27^. A recent mechanistic study demonstrated that the combination of lacosamide and phenytoin, which preferentially bind the slow- and fast-inactivated states of NaV channels, respectively, can produce supra-additive inhibition during prolonged depolarization, but infra-additive effects during intermediate depolarization^28^. These findings suggest that within-class NaV combinations may yield state-dependent effects and therefore require careful clinical monitoring.

Computational studies have become increasingly important in biomedical research. The recent application of machine learning models in healthcare has further expanded this field, creating new opportunities to map complex drug profiles to treat various disorders. To elucidate the complex process of drug-target interactions, it is essential to advance computational methodologies that integrate critical data from diverse sources^29–32^. This approach provides a robust foundation for understanding and mapping drug-target relationships. DRIFT is a deep learning model designed to map the targets of chemical compounds or drugs^33^. It uses a neural network architecture based on the YUEL algorithm to predict binding between a compound and its potential targets^34^. While understanding these individual interactions is critical, the context of pathways reveals their biological significance, as proteins function collectively within intricate networks. This network-based view is crucial because complex diseases often result from disruptions to multiple pathway components. By elucidating the specific relationships between drug targets and these disease-associated pathways, we can better understand disease mechanisms and pave the way for more efficacious therapies.

Given the critical gap in epilepsy therapies, we focused this study on selected NaV-targeting ASMs to investigate their off-target effects. Here, we present a computational pipeline that uses a neural network to elucidate the complex pharmacology of these ASMs. Our study first maps the protein targets for each ASM, including cenobamate, a more recently approved medication with a dual mechanism of action. We then expand this analysis to identify shared and unique targets across these drugs, providing crucial insights into their collective mechanisms and the molecular basis for rational polypharmacy.

## Methods

### Data Curation and Compilation

The initial dataset, comprising a selection of ASMs, was systematically compiled from the PubChem database. The specific compounds in this dataset, along with their corresponding SMILES (Simplified Molecular Input Line Entry System), are listed in Table 1. Eslicarbazepine is the (S)-enantiomer of licarbazepine, so licarbazepine was prepared as the (R)-enantiomer for the docking studies to keep the two compounds distinct.

**Table 1:**
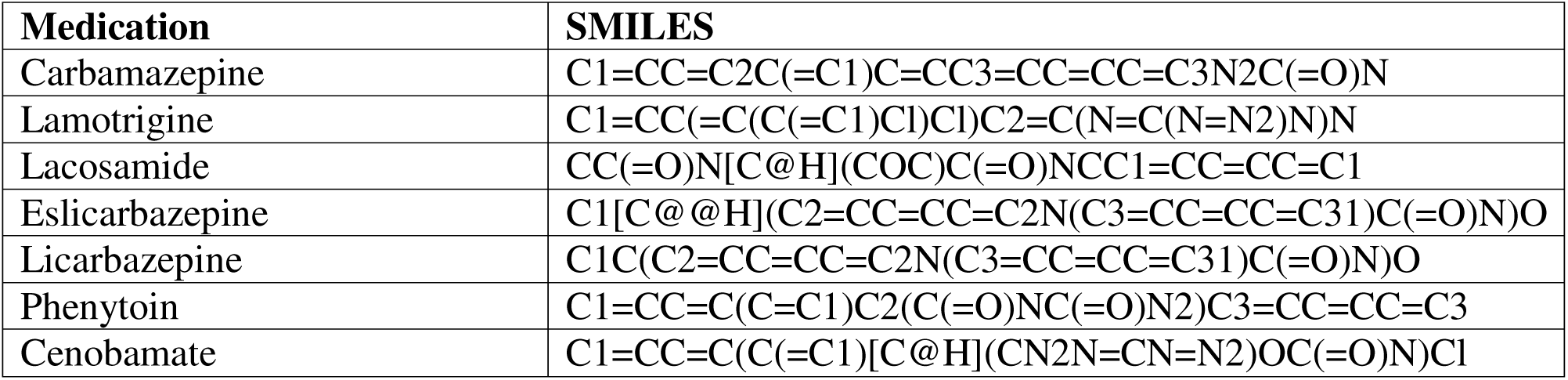
List of the sodium channel-targeting anti-seizure medications used in this study and their SMILES.

| **Medication** | **SMILES** |
|---|---|
| Carbamazepine | C1=CC=C2C(=C1)C=CC3=CC=CC=C3N2C(=O)N |
| Lamotrigine | C1=CC(=C(C(=C1)Cl)Cl)C2=C(N=C(N=N2)N)N |
| Lacosamide | CC(=O)N[C@H](COC)C(=O)NCC1=CC=CC=C1 |
| Eslicarbazepine | C1[C@@H](C2=CC=CC=C2N(C3=CC=CC=C31)C(=O)N)O |
| Licarbazepine | C1C(C2=CC=CC=C2N(C3=CC=CC=C31)C(=O)N)O |
| Phenytoin | C1=CC=C(C=C1)C2(C(=O)NC(=O)N2)C3=CC=CC=C3 |
| Cenobamate | C1=CC=C(C(=C1)[C@H](CN2N=CN=N2)OC(=O)N)Cl |

### Drug Target Identification (DRIFT)

DRIFT is a computational pipeline designed for proteome-wide identification of protein binders for bioactive small molecules^33^. It employs a multi-faceted approach, beginning with two-dimensional (2D) and three-dimensional (3D) chemical similarity searches to identify compounds structurally similar to a query molecule. YUEL, an attention-based neural network, is implemented within the DRIFT workflow to rank potential protein targets^34^. This process underpins the generation and Tanimoto coefficient-based comparison of pharmacophore and molecular fingerprints from the query compound against a curated reference database.

DRIFT was selected for three reasons. First, unlike structure- or docking-based target-prediction methods, DRIFT requires only the two-dimensional structure of a query compound. It can therefore screen the annotated human proteome without a high-resolution structure for every candidate target, which is essential for comparing off-targets across seven ASMs. Second, its target-ranking module, YUEL, is an attention-based neural network trained on curated compound–protein affinity data that generalizes to compound–protein pairs absent from the training set, partially mitigating the bias of purely similarity-based tools toward heavily studied drugs. Third, DRIFT has been benchmarked against established target-prediction tools and applied to structurally diverse ligands. The target predictions reported here were generated using the DRIFT web server. Importantly, DRIFT is intended here not to guide clinical prescribing directly but to nominate candidate off-targets and shared pathways that can be prioritized for experimental validation and, in the longer term, inform mechanism-guided hypotheses about ASM tolerability and combination therapy.

### Drug-Target Pathway Mapping

We mapped pathways associated with the identified protein targets using the Reactome database^35^. We first used an in-house Python script to map UniProt identifiers to their respective pathways^36^. For targets successfully mapped to UniProt IDs, the initial prediction scores were normalized to create a relative weighting system. The normalized score *w*_i_ for each target *i* was calculated by dividing its original score by the sum of all valid target scores in the dataset:

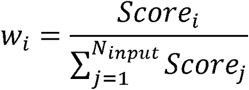

where *N_input_* is the total number of input genes successfully mapped. Here, the mapped entities are the predicted protein targets, each represented by its corresponding gene (via UniProt-to-gene mapping) for Reactome pathway analysis.

Pathway analysis was performed using the Reactome database. The UniProt-to-Reactome mapping was used to retrieve and filter annotations exclusively for *Homo sapiens*. A reference set containing all unique human genes represented in Reactome was established as the statistical background. The UniProt IDs mapped for the targets were then cross-referenced against the genes associated with each unique Reactome pathway to determine the number of mapped entities. To prioritize biologically relevant pathways, a custom pathway-scoring metric was calculated for each pathway that contained at least one mapped entity. The pathway score accounts for both the normalized weights of overlapping genes and the proportion of input genes mapped to that pathway. The score S for a pathway is defined as:

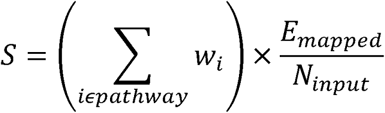

where *w_i_* is the normalized weight of the mapped gene, *E_mapped_* is the number of mapped entities in the specific pathway, and *N_input_* is the total number of mapped genes. Further, the raw pathway scores were subsequently scaled using Min-Max normalization to fall within a predefined range of 0.1 to 0.9. This systematic approach enabled us to predict pathways associated with the identified targets accurately and to enhance our understanding of the biological processes influenced by the compounds investigated.

### Drug-Target Similarity Analysis

To assess relationships between the drugs and their predicted targets, we used DRIFT-derived target profiles to generate vector representations for each ASM. These vectors served as the basis for computing cosine similarity scores, enabling the quantification of compound-target relationships. Pairwise cosine similarity values computed from drug-target vectors are visualized as a heatmap using matplotlib.

### Molecular Docking Studies

To further elucidate the atomistic interactions between the NaV-targeting ASMs and their predicted targets, we performed molecular docking using the MedusaDock program. Each drug was docked against its top-ranked target predicted by DRIFT. The initial conformations for computational studies of the protein targets were obtained from RCSB. The structures were subsequently optimized using the CHIRON program^37^, which efficiently minimizes steric clashes through short, discrete molecular dynamics simulations. The optimized structures were utilized for molecular docking studies and discrete molecular dynamics simulations. All structural visualizations were performed using UCSF ChimeraX^38^. MedusaDock has been extensively benchmarked, demonstrating rapid sampling efficiency and high prediction accuracy in computational docking studies^39–41^. Each drug was docked against the NaV subtype ranked highest for that drug by DRIFT, so that the docked structure matches the subtype reported in the target predictions. We therefore performed molecular docking of carbamazepine, eslicarbazepine, licarbazepine, lacosamide, and cenobamate against the NaV type IX (Nav1.7) alpha subunit (PDB 8S9B), lamotrigine against the NaV type II (Nav1.2) alpha subunit (PDB 6J8E), and phenytoin against the NaV type IV (Nav1.4) alpha subunit (PDB 6AGF). DRIFT returned no individual subtype for cenobamate, and Nav1.7 was used to match the other medications. The docking protocol targeted the central pore cavity of each channel. For Nav1.7 the cavity was defined by the position of the lacosamide molecule resolved in PDB 8S9B, and the equivalent cavity in Nav1.2 and Nav1.4 was located by sequence alignment of the alpha subunits. All non-protein molecules, including detergents and any bound ligand, were removed before docking. Each ligand underwent 100 independent docking iterations to ensure comprehensive sampling of possible binding conformations and enhance the reliability of the results. Binding energies were calculated using the MedusaDock scoring function. Poses were clustered, and the lowest-energy pose of the largest cluster is reported for each medication. Residues within 6.0 Å of the docked ligand pose were defined as interacting-site residues to identify all residues within the interaction sphere (Supplementary Table S1).

## Results

### Target identification and pathway mapping for the selected ASMs

We employed the DRIFT algorithm^33^ (Figure 1B) to identify proteome-wide targets for seven NaV-targeting ASMs (Table 1). The analysis revealed substantial variation in the predicted target landscape across drugs: eslicarbazepine and licarbazepine yielded the largest number of predicted targets (59 proteins each), followed by cenobamate (57), carbamazepine (52), phenytoin (51), lamotrigine (41), and lacosamide (18). The predicted targets were then mapped to biological pathways in the Reactome database.

### Carbamazepine and related dibenzazepine derivatives

Carbamazepine, licarbazepine, and eslicarbazepine represent a structurally related subgroup of NaV-targeting ASMs within the dibenzazepine carboxamide family. Although these compounds share a common tricyclic scaffold, differences in their metabolic and pharmacodynamic properties may contribute to distinct target and pathway landscapes^42^. We therefore considered these drugs together to assess how structural similarity relates to shared or divergent proteome-wide interactions.

For carbamazepine, DRIFT ranked the NaV alpha subunit family as the top target, consistent with its established action as a NaV blocker. The NaV type IX (Nav1.7) alpha subunit was the highest-ranked individual subtype. Furthermore, the model predicted interactions with P2X purinoceptor 4, organic anion transporters, UDP-glucuronosyltransferases, and cytochrome P450 enzymes, consistent with transport- and metabolism-related activity. Pathway mapping of these predicted targets revealed enrichment in Aspirin ADME, glucuronidation, and rapid depolarisation, indicating that carbamazepine may influence drug metabolism beyond simple NaV blockade (Figure 2A).

**Figure 2.**
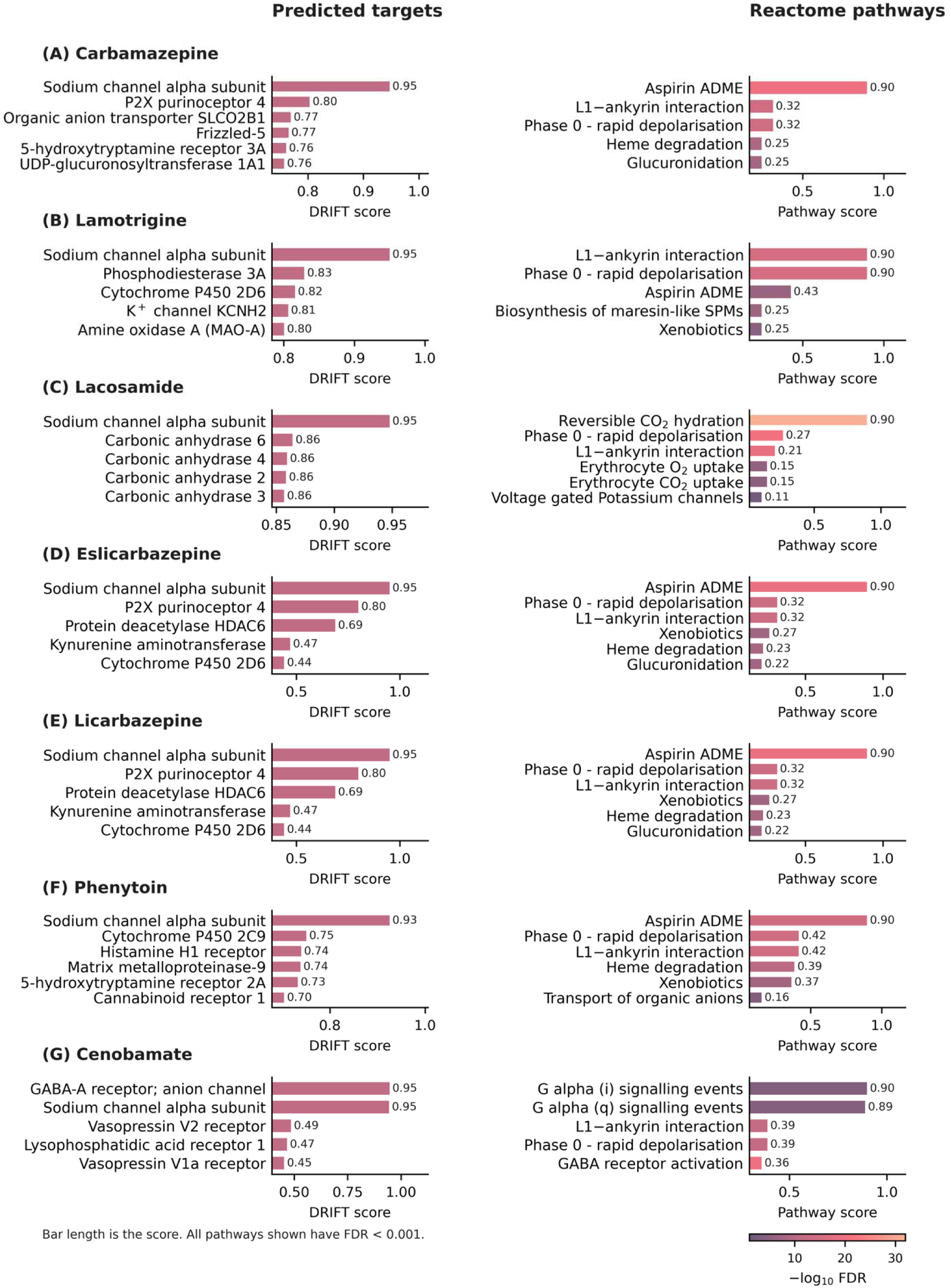
Proteome-wide target prediction and pathway analysis for selected anti-seizure medications. For each medication, the left subpanel shows the highest-ranked DRIFT-predicted protein targets and the right subpanel the most enriched Reactome pathways, both as bars whose length is the score. Pathway bars are coloured by false discovery rate, as −log10 FDR; every pathway shown has an FDR below 0.001. The complete lists are given in Supplementary Table S2 for the predicted targets and Supplementary Table S3 for the enriched pathways. (A) Carbamazepine displayed a sodium-channel-centered target profile with additional transporter- and metabolism-related proteins and pathway enrichment in aspirin ADME, glucuronidation, and rapid depolarisation. (B) Lamotrigine showed a broad target profile including voltage-gated sodium channel subunits, phosphodiesterase 3A, the potassium channel KCNH2, and transporter-related proteins, with enrichment in L1-ankyrin interactions, rapid depolarisation, and ADME-related pathways. (C) Lacosamide exhibited a relatively restricted target profile including the sodium channel alpha subunit family and the carbonic anhydrases, with strongest enrichment in carbon dioxide hydration and additional associations with rapid depolarisation and voltage-gated potassium channels. (D) Eslicarbazepine included the sodium channel alpha subunit family, P2X purinoceptor 4, protein deacetylase HDAC6, and cytochrome P450 enzymes, with enrichment in ADME-related pathways, xenobiotic metabolism, glucuronidation, and rapid depolarisation. (E) Licarbazepine returned predictions identical to eslicarbazepine, because the fingerprint used by DRIFT does not distinguish stereoisomers, with the same target set and the same enrichment in ADME-related pathways, xenobiotic metabolism, and glucuronidation. (F) Phenytoin included the sodium channel alpha subunit family, cytochrome P450 2C9, the histamine H1 receptor, and the cannabinoid CB1 receptor, with enrichment in aspirin ADME, organic anion transport, and xenobiotic metabolism. (G) Cenobamate included the GABA-A receptor and the sodium channel alpha subunit family, with enrichment in G alpha signalling events, GABA receptor activation, and rapid depolarisation.

For eslicarbazepine, DRIFT ranked the NaV alpha subunit family as the top target, consistent with published evidence of its activity at NaV channels. The model also predicted interactions with the P2X purinoceptor 4, protein deacetylase HDAC6, and cytochrome P450 enzymes, expanding the drug’s potential pharmacological profile. Mapping these targets to the Reactome database uncovered enrichment in ADME-related pathways, including Aspirin, Paracetamol, and Atorvastatin metabolism, together with xenobiotic metabolism and glucuronidation. These associations hint at downstream effects on drug transport networks and metabolic processing. Additionally, we detected enrichment in rapid depolarisation and the interaction between L1 and ankyrins, consistent with its action at NaV channels (Figure 2D).

For licarbazepine, DRIFT ranked the NaV alpha subunit family as the top target, consistent with its action as a NaV blocker. Furthermore, the model predicted interactions with the P2X purinoceptor 4, protein deacetylase HDAC6, and cytochrome P450 enzymes, consistent with transport- and metabolism-related activity. Because the fingerprint used by DRIFT does not distinguish stereoisomers, licarbazepine and eslicarbazepine returned identical target predictions. Pathway mapping of these predicted targets revealed enrichment in ADME-related pathways including Aspirin, Paracetamol, and Atorvastatin metabolism, indicating that licarbazepine may influence metabolic networks beyond primary NaV blockade (Figure 2E).

### Lamotrigine

Our analysis revealed a broad landscape of potential lamotrigine targets. The NaV alpha subunit family ranked highest. In addition, several individual NaV subunits (Nav1.1, Nav1.2, Nav1.3, Nav1.4, Nav1.5, Nav1.6, Nav1.7, and Nav1.8) were represented among the predicted targets. This broader predicted NaV-subunit profile may contribute to lamotrigine’s broad clinical activity across diverse seizure types. We also pinpointed the phosphodiesterase 3A and the potassium channel KCNH2 as candidates warranting experimental validation. At the pathway level, the predicted targets mapped to ankyrin-L1 interactions linked to NaV and rapid depolarisation. Additional enrichment in ADME-related pathways suggests that lamotrigine may also influence drug metabolism (Figure 2B).

### Lacosamide

For lacosamide, DRIFT highlighted the NaV alpha subunit family as the primary target, with the NaV type IX (Nav1.7) alpha subunit the highest-ranked individual subtype. This supports existing literature describing lacosamide as a modulator of slow inactivation in voltage-gated NaVs, although this effect is generally considered isoform-nonselective. The algorithm also captured the voltage-gated L-type calcium channel and the potassium ion channels KCND3 and KCNH2. While these predictions broaden lacosamide’s potential target landscape, their direct functional relevance requires experimental confirmation. Pathway mapping demonstrated that lacosamide possesses the most restricted profile among the analyzed drugs. Within this narrow scope, reversible hydration of carbon dioxide showed the greatest enrichment, consistent with prominent predictions for carbonic anhydrase targets. We also observed enrichment in Phase 0 rapid depolarisation, voltage-gated potassium channels, and interaction between L1 and Ankyrins, suggesting potential effects on neuronal excitability and cell adhesion signaling. Additionally, enrichment in erythrocyte carbon dioxide and oxygen exchange was observed, consistent with the prominent carbonic anhydrase predictions (Figure 2C).

### Phenytoin

For phenytoin, DRIFT ranked the NaV alpha subunit family as the top target, which aligns closely with phenytoin’s known mechanism, and the NaV type IV (Nav1.4) alpha subunit was the highest-ranked individual subtype. DRIFT also prioritized the cannabinoid CB1 receptor among the predicted targets. Because this computational prediction does not inherently establish phenytoin as a direct CB1 ligand, it necessitates future experimental validation. Furthermore, the model predicted cytochrome P450 2C9, the histamine H1 receptor, and matrix metalloproteinase 9. Pathway enrichment analysis highlighted Aspirin ADME and organic anion transport, reflecting phenytoin’s known susceptibility to metabolism- and transporter-mediated drug interactions (Figure 2F).

### Cenobamate

For cenobamate, DRIFT ranked the GABA-A receptor and the NaV alpha subunit family as the two highest targets, with near-identical scores, consistent with its recognized dual mechanism. The model also predicted interactions with the vasopressin V2 and V1a receptors, the lysophosphatidic acid receptors, and caspase-3, expanding the drug’s potential pharmacological profile. Pathway mapping of these predicted targets revealed enrichment in G alpha (i) and G alpha (q) signalling events, GABA receptor activation, and rapid depolarisation, consistent with a mechanism that combines sodium channel modulation with enhancement of inhibitory signaling (Figure 2G).

Comparing the predicted target profiles across these seven sodium-channel-targeting ASMs revealed both shared and distinct pharmacological features. All seven medications shared the NaV alpha subunit family as a top-ranked target, mirroring their shared inactivation-related mechanisms. Eslicarbazepine and licarbazepine displayed the broadest target landscapes, in contrast to lacosamide’s more restricted predicted profile, reflecting their respective broad-spectrum and selective clinical utilities. At the pathway level, we observed recurrent enrichment in rapid depolarisation and the interaction between L1 and ankyrins across all seven drugs. These shared pathway signatures imply convergent effects on neuronal excitability, alongside drug-metabolism pathways that extend well beyond traditional NaV blockade.

### Target similarity profiling

To quantify the overall target profile similarity among the seven ASMs, we developed a computational workflow that uses DRIFT predictions, target vectorization, and cosine similarity (Figure 3). The resulting target similarity matrix demonstrates distinct relationships among the evaluated drugs. Carbamazepine, eslicarbazepine, and licarbazepine showed the highest pairwise similarity, suggesting overlapping predicted target profiles and shared off-target landscapes, whereas lamotrigine and phenytoin formed a second, weaker pairing. In contrast, lacosamide emerges as a distinct outlier within this group. The matrix shows the lowest similarity values between lacosamide and the other ASMs, indicating that its predicted proteome-wide target landscape is highly dissimilar to those of the other evaluated NaV modulators, with cenobamate the next most distinct.

**Figure 3.**
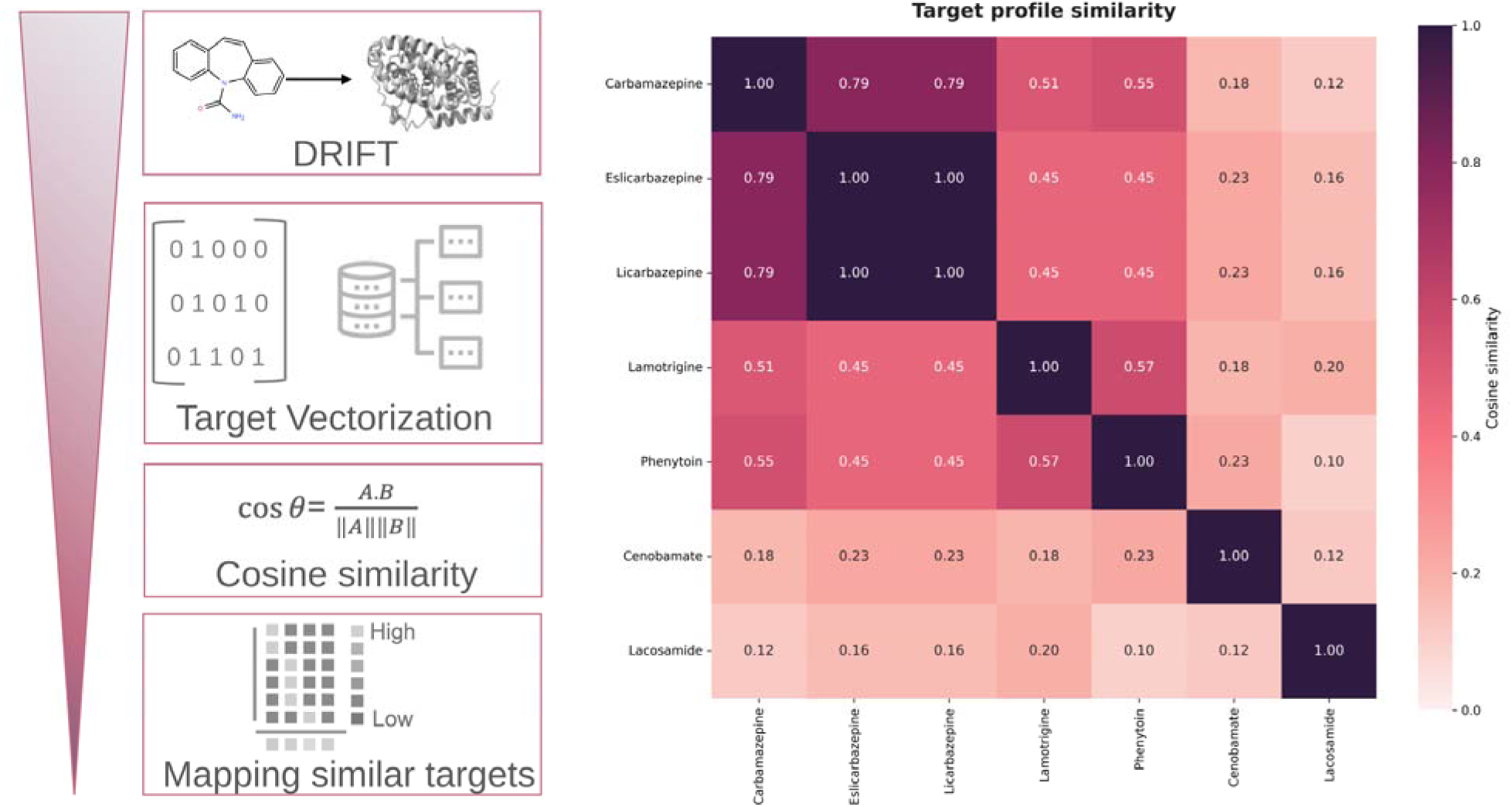
Comparative target similarity analysis across seven ASMs. Schematic representation of the workflow used for target vectorization and cosine similarity-based comparison of DRIFT-predicted target profiles (left). Heatmap of the pairwise cosine similarity matrix, showing relative clustering of carbamazepine, eslicarbazepine, and licarbazepine, and of lamotrigine with phenytoin, with lacosamide exhibiting a distinct and comparatively dissimilar proteome-wide target landscape and cenobamate the next most distinct (right). Values range from 0 to 1.

## Molecular Docking Studies

To further elucidate the atomistic interactions of the NaV ASMs with their predicted targets, we performed molecular docking using the MedusaDock program. Each drug was docked against the NaV subtype ranked highest for that drug by DRIFT (Table 2). The structural models illustrate the NaV alpha subunits (Types II, IV, and IX) in green, with the docked ASMs (yellow) occupying the central pore cavity of each channel (Figure 4). Carbamazepine docked within the NaV type IX (Nav1.7) alpha subunit with a docking score of −27.32 kcal/mol, showing proximity to Leu398, Ala402, Leu964, and Ile1457 (Figure 4A). In comparison, lamotrigine docked within the NaV type II (Nav1.2) alpha subunit with a score of −23.91 kcal/mol, forming interactions with Leu421, Ala425, Leu982, and Leu1776 (Figure 4B). Eslicarbazepine docked within the NaV type IX (Nav1.7) alpha subunit showed a score of −27.76 kcal/mol, interacting closely with a binding site containing residues Leu398, Ala402, Leu967, and Leu1760 (Figure 4D). Similarly, lacosamide docked into the same subunit with a score of −27.37 kcal/mol, interacting with Leu964, Ile1457, Asn1461, and Leu1760 (Figure 4C). These poses identify residues lining the predicted binding site. Phenytoin yielded the highest docking score (−30.75 kcal/mol) within the NaV type IV (Nav1.4) alpha subunit (Figure 4F), with the pose surrounded by Leu443, Leu798, Ile1291, and Ile1594. Furthermore, licarbazepine docked within the NaV type IX (Nav1.7) alpha subunit showed a binding score of −26.74 kcal/mol, interacting closely with binding site residues Leu964, Leu967, Ile1453, and Ile1457 (Figure 4E), and cenobamate docked into the same subunit with a score of −28.53 kcal/mol, contacting Glu406, Leu964, Leu967, and Ile1756 (Figure 4G). However, this interesting finding still requires empirical validation to substantiate our computational predictions further. In every case the pose occupied the central pore cavity and contacted residues drawn from all four homologous repeats of the channel, and the contacted positions were equivalent across the three subtypes examined. More generally, the dense packing of specific hydrophobic and polar residues, along with the docking scores, suggests that these NaV-blocking ASMs could form stable interactions within their predicted binding sites. Hence, these docking results should be viewed as structural support for the DRIFT predictions and could help generate hypotheses for these off-targets, which require further experimental validation.

**Figure 4.**
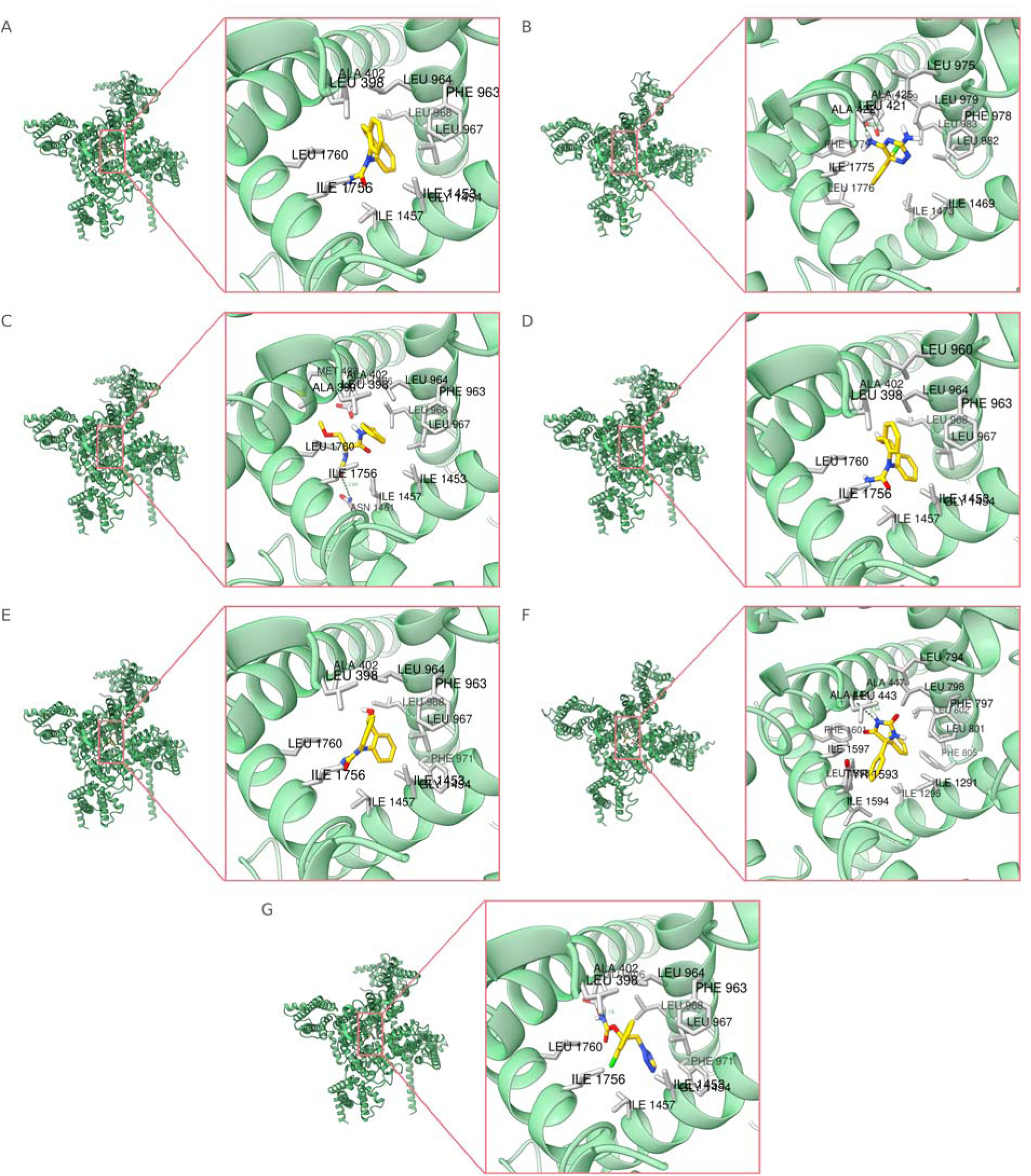
Structural models of docked anti-seizure medications in the central pore cavity of their top-ranked sodium channel subtype. (A) Carbamazepine, (C) lacosamide, (D) eslicarbazepine, (E) licarbazepine, and (G) cenobamate are shown docked in the NaV type IX alpha subunit, (B) lamotrigine in the NaV type II alpha subunit, and (F) phenytoin in the NaV type IV alpha subunit. The channel is shown in green and the medication in yellow. Insets show representative residues within 4.5 Å of each ligand pose, labeled with the residue numbering of the deposited structure, providing structural context for the predicted interactions obtained from MedusaDock analysis. Reported docking scores are listed in Table 2.

**Table 2:**
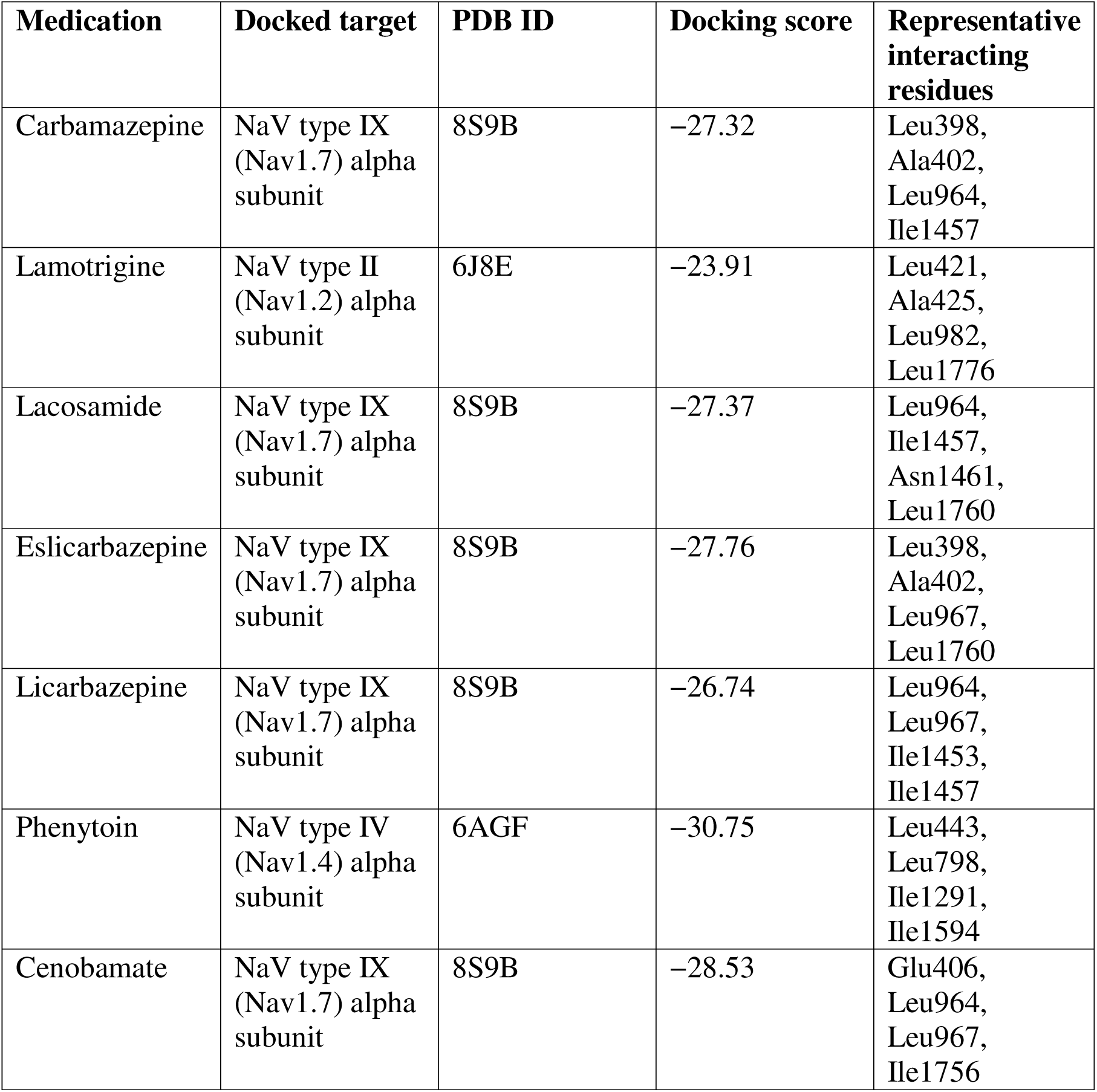
Molecular docking of the sodium channel subtype predicted by DRIFT for each medication. All scores are reported in kcal/mol. Scores are the MedusaDock energy of the lowest-energy pose of the largest cluster from 100 independent docking runs.

| **Medication** | **Docked target** | **PDB ID** | **Docking score** | **Representative interacting residues** |
|---|---|---|---|---|
| Carbamazepine | NaV type IX (Nav1.7) alpha subunit | 8S9B | −27.32 | Leu398, Ala402, Leu964, Ile1457 |
| Lamotrigine | NaV type II (Nav1.2) alpha subunit | 6J8E | −23.91 | Leu421, Ala425, Leu982, Leu1776 |
| Lacosamide | NaV type IX (Nav1.7) alpha subunit | 8S9B | −27.37 | Leu964, Ile1457, Asn1461, Leu1760 |
| Eslicarbazepine | NaV type IX (Nav1.7) alpha subunit | 8S9B | −27.76 | Leu398, Ala402, Leu967, Leu1760 |
| Licarbazepine | NaV type IX (Nav1.7) alpha subunit | 8S9B | −26.74 | Leu964, Leu967, Ile1453, Ile1457 |
| Phenytoin | NaV type IV (Nav1.4) alpha subunit | 6AGF | −30.75 | Leu443, Leu798, Ile1291, Ile1594 |
| Cenobamate | NaV type IX (Nav1.7) alpha subunit | 8S9B | −28.53 | Glu406, Leu964, Leu967, Ile1756 |

## Discussion

Our systems-level analysis shows that, while these seven ASMs primarily act as NaV blockers, their divergent proteome-wide target landscapes may contribute to differences in their clinical profiles. Beyond classical NaV blockade, we found that the medications share recurrent enrichment in drug-metabolism and transporter pathways, suggesting overlapping routes of elimination. Furthermore, our computational pipeline effectively generates novel, testable hypotheses, such as the predicted interaction between phenytoin and the cannabinoid CB1 receptor.

Viewed together, these results reframe sodium-channel–targeting ASMs not as simple NaV blockers but as multi-target, multi-pathway drugs, whose broader interactomes can and probably do shape clinical outcomes. That shift has several direct implications for epilepsy treatment. Drugs we often lump together, such as carbamazepine, lamotrigine, lacosamide, phenytoin, eslicarbazepine, licarbazepine, and cenobamate, actually have distinct proteome-wide target landscapes. For instance, lamotrigine showed a broad landscape, targeting a wide range of NaV subtypes, phosphodiesterase 3A, and the potassium channel KCNH2, whereas lacosamide showed a narrow, carbonic anhydrase–dominant profile. Phenytoin yielded an unexpected prediction at the cannabinoid CB1 receptor, while the structurally related dibenzazepines carbamazepine, eslicarbazepine, and licarbazepine shared strong overlap in organic-anion transporters and ADME pathways.

These distinct interactomes offer a conceptual framework for several long-standing clinical observations: why some NaV-targeting ASMs are effective for focal but not generalized seizures, why certain combinations appear synergistic whereas others increase toxicity, and why individual patients may respond to one NaV blocker but not another. Conceptually, this shifts the field from viewing NaV blockers as interchangeable to recognizing that each NaV-targeting ASM has a unique multi-target signature that shapes efficacy, tolerability, and optimal combinations.

The target-similarity analysis is consistent with this view. Carbamazepine, eslicarbazepine, and licarbazepine, and, to a lesser extent, phenytoin and lamotrigine, share targets enriched in aspirin and atorvastatin metabolism and organic-anion transport, suggesting overlapping toxicity and drug–drug interaction pathways when such agents are combined. Lacosamide was the clear outlier, showing the lowest average similarity with the other NaV ASMs and therefore the most distinct predicted target landscape; this divergence provides a mechanistic rationale for its frequent clinical use as an add-on therapy. Antiseizure medications are commonly combined to treat seizures refractory to monotherapy, and current practice guidelines from the American Academy of Neurology and the American Epilepsy Society list a series of drugs that can be added to existing therapies^43^. However, they do not recommend any specific drug combinations that are likely to be efficacious. A rigorous combination drug therapy trial would be challenging to design and cost-prohibitive. The current computational approach mapping shared and divergent interactomes may help prioritize rational pairings that would be too costly to establish empirically in a large prospective combination trial.

The pathway enrichments reported here warrant careful interpretation, because several, such as “Aspirin ADME” and organic-anion transport, are not proposed as antiseizure targets. These are Reactome annotations for shared drug-metabolizing enzymes (e.g., UDP-glucuronosyltransferases and cytochrome P450s) and membrane transporters that process many co-administered medications. Their recurrent enrichment therefore marks where NaV-targeting ASMs are likely to share metabolic and transporter handling, and thus where pharmacokinetic drug–drug interactions and overlapping dose-limiting toxicities are most probable during polytherapy. These pathway signatures translate into concrete, testable expectations about tolerability and interaction risk, rather than direct antiseizure mechanisms.

Several caveats apply to all of the predictions reported here. DRIFT identifies candidate targets by chemical-similarity inference and neural-network ranking, such as phenytoin at the CB1 receptor, which indicates a hypothesis worth testing rather than an established direct interaction. Because DRIFT infers targets from chemical similarity, predicted target counts (e.g., 59 for eslicarbazepine vs. 18 for lacosamide) reflect database coverage as much as true target breadth and should be read as relative, not absolute. Experimental screening is needed to confirm which predicted off-targets are genuinely engaged. Likewise, the docking scores provide structural plausibility for the predicted poses but are not a substitute for binding or functional assays. Accordingly, the off-targets and pathway associations described for every drug in this study require experimental validation before clinical inferences are drawn.

Because DRIFT ranks candidate targets from chemical similarity to annotated compounds and neural-network-predicted affinity, it preferentially recovers interactions well represented as compound–protein affinity records and can miss physiologically important interactions that are state- or use-dependent or mediated by mechanisms absent from those databases. Consistent with this, DRIFT did not recover several experimentally established interactions of the drugs studied here. Lamotrigine augments the hyperpolarization-activated cyclic-nucleotide-gated (HCN) current (Ih) that contributes to its dendritic and anti-absence activity^44^, and eslicarbazepine inhibits CaV3.2 (T-type) calcium channels^45^; neither interaction appeared among the top-ranked predictions. These false negatives indicate that the predicted interactomes are incomplete and should be read as a complement to, not a substitute for, experimental pharmacology.

The model does not incorporate tissue expression or physiological context, which can yield predictions difficult to reconcile with known neuropharmacology. Carbamazepine’s highest-ranked individual sodium channel was Nav1.7 (type IX), which is enriched in peripheral sensory neurons, whereas the neuronally enriched Nav1.6 (type VIII) would be the expected target for an ASM.^46^ Likewise, phenytoin’s highest-ranked individual subtype was Nav1.4 (type IV), which is expressed predominantly in skeletal muscle. Because ranking reflects chemical similarity-weighted binding rather than where a channel is expressed or whether its engagement is physiologically consequential, isoform- and tissue-level specificity is not captured. High-ranking non-neuronal predictions, such as CB1 for phenytoin, should be treated as chemically plausible hypotheses for experimental testing rather than as validated neuronal mechanisms.

Predicted target range also does not map directly onto the clinical spectrum. Lamotrigine was predicted to engage Nav1.7 (type IX), a channel central to nociception, yet it lacks the analgesic efficacy well established for carbamazepine. This discordance reinforces the idea that a predicted binding interaction need not translate into a functional or clinically meaningful effect.

The target-similarity analysis (Figure 3) is subject to the same constraints. Eslicarbazepine and lacosamide did not show strong target-profile similarity even though both enhance slow inactivation of voltage-gated sodium channels^24^. Because cosine similarity is computed over the entire predicted target vector rather than over a shared functional mechanism, two drugs that converge on the same channel state through different chemotypes can appear dissimilar; the matrix should therefore be read as reflecting global chemical-target overlap, not mechanistic equivalence.

## Conclusion

In this study, we used a computational approach to characterize proteome-wide targets and pathways for seven ASMs that modulate NaVs. Our results suggest that these NaV-blocking ASMs exhibit distinct interaction profiles. Further, our study identified candidate off-targets, including the cannabinoid CB1 receptor for phenytoin and the carbonic anhydrases for lacosamide, which may serve as hypotheses for future experimental validation. Overall, our computational framework integrates known targets with computationally predicted off-targets, which could inform future research directions and the rational design of efficacious therapeutic strategies for treating epilepsy.

## Author contributions

S.E. formal analysis; investigation; methodology (development of data analysis procedures); writing—original draft. E.A. conceptualization; writing—review & editing. J.K., H.G., and N.D. Conceptualization; resources; supervision; writing—review & editing.

## Acknowledgment

N.V.D. acknowledges support from the National Institutes of Health (R35 GM134864 and R01 AT012053-01)

## Conflict of interest statement

None of the authors has any conflict of interest to disclose.

## Ethics publication statement

We confirm that we have read the Journal’s position on ethical publication issues and affirm that this study is consistent with those guidelines.

## Data availability statement

The data reported in this article will be made available upon reasonable request.

## Notes

### Competing Interest Statement

The authors have declared no competing interest.

