## Supplementary tables for "Elucidating the Sodium Channel-Targeting Anti-Seizure Medication Interactome: A Drug-Target Pathway Analysis"

Supplementary Table S1: All residues within 6.0 Å of the docked pose of each medication.

| **Medication** | **Number of residues** | **Residues within 6.0 Å** |
| --- | --- | --- |
| Carbamazepine | 21 | Leu398, Ala399, Ala402, Glu406, Leu960, Phe963, Leu964, Leu967, Leu968, Phe971, Ser972, Ile1453, Gly1454, Ile1457, Asp1458, Asn1461, Val1752, Asn1753, Ile1756, Ala1757, Leu1760 |
| Lamotrigine | 23 | Asn418, Ile420, Leu421, Ala422, Val423, Val424, Ala425, Met426, Glu429, Leu975, Phe978, Leu979, Leu982, Leu983, Phe986, Ile1469, Ile1473, Asn1477, Tyr1771, Ile1772, Ile1775, Leu1776, Phe1779 |
| Lacosamide | 18 | Leu398, Ala399, Ala402, Met403, Glu406, Phe963, Leu964, Leu967, Leu968, Ile1453, Ile1457, Asn1461, Ile1756, Ala1757, Leu1760, Glu1761, Phe1763, Ser1764 |
| Eslicarbazepine | 21 | Leu398, Ala399, Ala402, Met403, Glu406, Leu960, Phe963, Leu964, Leu967, Leu968, Phe971, Ile1453, Gly1454, Val1455, Ile1457, Asp1458, Asn1461, Asn1753, Ile1756, Ala1757, Leu1760 |
| Licarbazepine | 19 | Leu398, Ala402, Met403, Tyr405, Glu406, Phe963, Leu964, Leu967, Leu968, Phe971, Ile1453, Gly1454, Val1455, Ile1457, Asp1458, Asn1461, Ile1756, Ala1757, Leu1760 |
| Phenytoin | 24 | Asn440, Ile442, Leu443, Ala444, Val445, Val446, Ala447, Met448, Leu794, Phe797, Leu798, Leu801, Leu802, Phe805, Ile1291, Gly1292, Ile1295, Val1590, Tyr1593, Ile1594, Ala1595, Ile1597, Leu1598, Phe1601 |
| Cenobamate | 21 | Leu398, Ala399, Val401, Ala402, Met403, Glu406, Phe963, Leu964, Leu967, Leu968, Phe971, Asn1450, Ile1453, Gly1454, Val1455, Ile1457, Asp1458, Asn1461, Ile1756, Ala1757, Leu1760 |

Supplementary Table S2A: All 52 protein targets predicted by DRIFT for carbamazepine, ranked by DRIFT score. Figure 2A shows only the highest-ranked of these.

| **Rank** | **Predicted target** | **DRIFT score** |
| --- | --- | --- |
| 1 | Sodium channel alpha subunit | 0.948 |
| 2 | P2X purinoceptor 4 | 0.804 |
| 3 | Solute carrier organic anion transporter family member 2B1 | 0.770 |
| 4 | Frizzled-5 | 0.766 |
| 5 | 5-hydroxytryptamine receptor 3A | 0.761 |
| 6 | Cytochrome P450 2C19 | 0.761 |
| 7 | UDP-glucuronosyltransferase 1A6 | 0.759 |
| 8 | UDP-glucuronosyltransferase 1A1 | 0.758 |
| 9 | Cytochrome P450 2B6 | 0.754 |
| 10 | UDP-glucuronosyltransferase 1A4 | 0.753 |
| 11 | Acetylcholinesterase | 0.749 |
| 12 | Cytochrome P450 3A4 | 0.747 |
| 13 | Glucocorticoid receptor | 0.746 |
| 14 | Cytochrome P450 2D6 | 0.744 |
| 15 | Androgen receptor | 0.743 |
| 16 | UDP-glucuronosyltransferase 1A3 | 0.742 |
| 17 | Cytochrome P450 2C9 | 0.742 |
| 18 | Albumin | 0.738 |
| 19 | Solute carrier family 22 member 1 | 0.737 |
| 20 | Frizzled-7 | 0.735 |
| 21 | Solute carrier organic anion transporter family member 1B1 | 0.734 |
| 22 | UDP-glucuronosyltransferase 2B7 | 0.734 |
| 23 | UDP-glucuronosyltransferase 1A9 | 0.733 |
| 24 | Cytochrome P450 1A2 | 0.731 |
| 25 | Deoxynucleoside triphosphate triphosphohydrolase SAMHD1 | 0.728 |
| 26 | Sodium channel protein type 9 subunit alpha | 0.724 |
| 27 | Cytochrome P450 2C8 | 0.724 |
| 28 | Solute carrier organic anion transporter family member 1B3 | 0.721 |
| 29 | Chymotrypsin-C | 0.720 |
| 30 | Tyrosyl-DNA phosphodiesterase 1 | 0.718 |
| 31 | Sodium channel protein type 5 subunit alpha | 0.707 |
| 32 | Sodium channel protein type 4 subunit alpha | 0.705 |
| 33 | Malate dehydrogenase, cytoplasmic | 0.704 |
| 34 | Protein deacetylase HDAC6 | 0.690 |
| 35 | Sodium channel protein type 2 subunit alpha | 0.690 |
| 36 | ATP-dependent translocase ABCB1 | 0.690 |
| 37 | ATP-binding cassette sub-family C member 2 | 0.690 |
| 38 | P2X purinoceptor 3 | 0.442 |
| 39 | P2X purinoceptor 1 | 0.431 |
| 40 | Melatonin receptor type 1B | 0.406 |
| 41 | P2X purinoceptor 7 | 0.401 |
| 42 | Melatonin receptor type 1A | 0.392 |
| 43 | Survival motor neuron protein | 0.384 |
| 44 | Glucagon-like peptide 1 receptor | 0.359 |
| 45 | P2X purinoceptor 2 | 0.358 |
| 46 | Polypyrimidine tract-binding protein 1 | 0.358 |
| 47 | NAD-dependent protein deacetylase sirtuin-1 | 0.352 |
| 48 | Ubiquitin carboxyl-terminal hydrolase 2 | 0.350 |
| 49 | NAD-dependent protein deacetylase sirtuin-2 | 0.339 |
| 50 | Cholinesterase | 0.337 |
| 51 | Serine/threonine-protein kinase PLK1 | 0.240 |
| 52 | Vitamin D3 receptor | 0.223 |

Supplementary Table S2B: All 41 protein targets predicted by DRIFT for lamotrigine, ranked by DRIFT score. Figure 2B shows only the highest-ranked of these.

| **Rank** | **Predicted target** | **DRIFT score** |
| --- | --- | --- |
| 1 | Sodium channel alpha subunit | 0.951 |
| 2 | cGMP-inhibited 3',5'-cyclic phosphodiesterase 3A | 0.830 |
| 3 | Cytochrome P450 2D6 | 0.817 |
| 4 | Voltage-gated inwardly rectifying potassium channel KCNH2 | 0.807 |
| 5 | Amine oxidase [flavin-containing] A | 0.801 |
| 6 | Multidrug and toxin extrusion protein 1 | 0.772 |
| 7 | Cytochrome P450 2C19 | 0.771 |
| 8 | Alpha-1A adrenergic receptor | 0.769 |
| 9 | Chymotrypsin-C | 0.768 |
| 10 | Cytochrome P450 3A4 | 0.766 |
| 11 | Cytochrome P450 1A2 | 0.761 |
| 12 | Cytochrome P450 2C9 | 0.757 |
| 13 | A-type voltage-gated potassium channel KCND3 | 0.755 |
| 14 | Solute carrier organic anion transporter family member 1B1 | 0.755 |
| 15 | Solute carrier organic anion transporter family member 1B3 | 0.752 |
| 16 | Aquaporin-4 | 0.750 |
| 17 | Solute carrier organic anion transporter family member 2B1 | 0.750 |
| 18 | Deoxynucleoside triphosphate triphosphohydrolase SAMHD1 | 0.748 |
| 19 | Multidrug and toxin extrusion protein 2 | 0.747 |
| 20 | UDP-glucuronosyltransferase 1A4 | 0.747 |
| 21 | Sodium channel protein type 2 subunit alpha | 0.741 |
| 22 | Sodium channel protein type 4 subunit alpha | 0.730 |
| 23 | Sodium channel protein type 9 subunit alpha | 0.730 |
| 24 | Sodium channel protein type 3 subunit alpha | 0.730 |
| 25 | Malate dehydrogenase, cytoplasmic | 0.729 |
| 26 | Albumin | 0.722 |
| 27 | Sodium channel protein type 5 subunit alpha | 0.711 |
| 28 | Tyrosine-protein phosphatase non-receptor type 11 | 0.697 |
| 29 | Sodium channel protein type 10 subunit alpha | 0.691 |
| 30 | Sodium channel protein type 1 subunit alpha | 0.690 |
| 31 | Protein deacetylase HDAC6 | 0.690 |
| 32 | Sodium channel protein type 8 subunit alpha | 0.690 |
| 33 | Cannabinoid receptor 2 | 0.503 |
| 34 | Oxidized purine nucleoside triphosphate hydrolase | 0.462 |
| 35 | Palmitoleoyl-protein carboxylesterase NOTUM | 0.390 |
| 36 | Casein kinase I isoform alpha | 0.337 |
| 37 | Dual specificity protein kinase CLK4 | 0.337 |
| 38 | Solute carrier family 2, facilitated glucose transporter member 1 | 0.315 |
| 39 | Dihydrofolate reductase | 0.285 |
| 40 | Aromatase | 0.258 |
| 41 | Nuclear receptor subfamily 1 group I member 2 | 0.257 |

Supplementary Table S2C: All 18 protein targets predicted by DRIFT for lacosamide, ranked by DRIFT score. Figure 2C shows only the highest-ranked of these.

| **Rank** | **Predicted target** | **DRIFT score** |
| --- | --- | --- |
| 1 | Sodium channel alpha subunit | 0.948 |
| 2 | Carbonic anhydrase 6 | 0.864 |
| 3 | Carbonic anhydrase 4 | 0.860 |
| 4 | Carbonic anhydrase 2 | 0.859 |
| 5 | Carbonic anhydrase 3 | 0.857 |
| 6 | Carbonic anhydrase 5B, mitochondrial | 0.856 |
| 7 | Carbonic anhydrase 14 | 0.852 |
| 8 | Carbonic anhydrase 1 | 0.848 |
| 9 | Carbonic anhydrase 9 | 0.844 |
| 10 | Carbonic anhydrase 13 | 0.827 |
| 11 | Carbonic anhydrase 12 | 0.825 |
| 12 | Sodium channel protein type 9 subunit alpha | 0.815 |
| 13 | Carbonic anhydrase 5A, mitochondrial | 0.812 |
| 14 | Carbonic anhydrase 7 | 0.812 |
| 15 | A-type voltage-gated potassium channel KCND3 | 0.738 |
| 16 | Protein deacetylase HDAC6 | 0.690 |
| 17 | Voltage-dependent L-type calcium channel subunit alpha-1C | 0.687 |
| 18 | Voltage-gated inwardly rectifying potassium channel KCNH2 | 0.671 |

Supplementary Table S2D: All 59 protein targets predicted by DRIFT for eslicarbazepine, ranked by DRIFT score. Figure 2D shows only the highest-ranked of these.

| **Rank** | **Predicted target** | **DRIFT score** |
| --- | --- | --- |
| 1 | Sodium channel alpha subunit | 0.953 |
| 2 | P2X purinoceptor 4 | 0.802 |
| 3 | Protein deacetylase HDAC6 | 0.690 |
| 4 | Kynurenine/alpha-aminoadipate aminotransferase, mitochondrial | 0.472 |
| 5 | Cytochrome P450 2D6 | 0.443 |
| 6 | Cytochrome P450 2C19 | 0.443 |
| 7 | Cytochrome P450 3A4 | 0.442 |
| 8 | Cytochrome P450 2C9 | 0.441 |
| 9 | Acetylcholinesterase | 0.436 |
| 10 | Tyrosyl-DNA phosphodiesterase 1 | 0.430 |
| 11 | Cytochrome P450 1A2 | 0.426 |
| 12 | Androgen receptor | 0.422 |
| 13 | 5-hydroxytryptamine receptor 3A | 0.419 |
| 14 | NAD-dependent protein deacetylase sirtuin-1 | 0.406 |
| 15 | Glucocorticoid receptor | 0.404 |
| 16 | NAD-dependent protein deacetylase sirtuin-2 | 0.400 |
| 17 | Aldehyde dehydrogenase 1A1 | 0.398 |
| 18 | Huntingtin | 0.365 |
| 19 | Epoxide hydrolase 1 | 0.350 |
| 20 | Solute carrier organic anion transporter family member 1B1 | 0.345 |
| 21 | Solute carrier family 22 member 12 | 0.339 |
| 22 | Cholinesterase | 0.322 |
| 23 | Deoxynucleoside triphosphate triphosphohydrolase SAMHD1 | 0.321 |
| 24 | Microphthalmia-associated transcription factor | 0.313 |
| 25 | Ribosomal protein S6 kinase beta-1 | 0.311 |
| 26 | Solute carrier organic anion transporter family member 1B3 | 0.307 |
| 27 | Aurora kinase A | 0.302 |
| 28 | Kynurenine--oxoglutarate transaminase 3 | 0.301 |
| 29 | UDP-glucuronosyltransferase 1A6 | 0.297 |
| 30 | Palmitoleoyl-protein carboxylesterase NOTUM | 0.295 |
| 31 | Nuclear factor erythroid 2-related factor 2 | 0.295 |
| 32 | Frizzled-5 | 0.295 |
| 33 | Kynurenine--oxoglutarate transaminase 1 | 0.294 |
| 34 | Frizzled-7 | 0.293 |
| 35 | Solute carrier family 22 member 1 | 0.292 |
| 36 | UDP-glucuronosyltransferase 2B7 | 0.291 |
| 37 | Vitamin D3 receptor | 0.288 |
| 38 | Aspartate aminotransferase, cytoplasmic | 0.283 |
| 39 | UDP-glucuronosyltransferase 1A4 | 0.283 |
| 40 | UDP-glucuronosyltransferase 1A3 | 0.282 |
| 41 | Cytochrome P450 2C8 | 0.281 |
| 42 | Cytochrome P450 2B6 | 0.279 |
| 43 | Chymotrypsin-C | 0.279 |
| 44 | Solute carrier organic anion transporter family member 2B1 | 0.277 |
| 45 | UDP-glucuronosyltransferase 1A1 | 0.276 |
| 46 | Albumin | 0.272 |
| 47 | UDP-glucuronosyltransferase 1A9 | 0.270 |
| 48 | Malate dehydrogenase, cytoplasmic | 0.251 |
| 49 | ATP-binding cassette sub-family C member 2 | 0.250 |
| 50 | Serine/threonine-protein kinase PLK1 | 0.247 |
| 51 | Solute carrier family 2, facilitated glucose transporter member 1 | 0.239 |
| 52 | TAR DNA-binding protein 43 | 0.239 |
| 53 | Solute carrier family 22 member 3 | 0.234 |
| 54 | Sodium channel protein type 9 subunit alpha | 0.232 |
| 55 | Sodium channel protein type 5 subunit alpha | 0.218 |
| 56 | Sodium channel protein type 4 subunit alpha | 0.215 |
| 57 | ATPase family AAA domain-containing protein 5 | 0.210 |
| 58 | Sodium channel protein type 2 subunit alpha | 0.204 |
| 59 | ATP-dependent translocase ABCB1 | 0.204 |

Supplementary Table S2E: All 59 protein targets predicted by DRIFT for licarbazepine, ranked by DRIFT score. Figure 2E shows only the highest-ranked of these.

| **Rank** | **Predicted target** | **DRIFT score** |
| --- | --- | --- |
| 1 | Sodium channel alpha subunit | 0.953 |
| 2 | P2X purinoceptor 4 | 0.802 |
| 3 | Protein deacetylase HDAC6 | 0.690 |
| 4 | Kynurenine/alpha-aminoadipate aminotransferase, mitochondrial | 0.472 |
| 5 | Cytochrome P450 2D6 | 0.443 |
| 6 | Cytochrome P450 2C19 | 0.443 |
| 7 | Cytochrome P450 3A4 | 0.442 |
| 8 | Cytochrome P450 2C9 | 0.441 |
| 9 | Acetylcholinesterase | 0.436 |
| 10 | Tyrosyl-DNA phosphodiesterase 1 | 0.430 |
| 11 | Cytochrome P450 1A2 | 0.426 |
| 12 | Androgen receptor | 0.422 |
| 13 | 5-hydroxytryptamine receptor 3A | 0.419 |
| 14 | NAD-dependent protein deacetylase sirtuin-1 | 0.406 |
| 15 | Glucocorticoid receptor | 0.404 |
| 16 | NAD-dependent protein deacetylase sirtuin-2 | 0.400 |
| 17 | Aldehyde dehydrogenase 1A1 | 0.398 |
| 18 | Huntingtin | 0.365 |
| 19 | Epoxide hydrolase 1 | 0.350 |
| 20 | Solute carrier organic anion transporter family member 1B1 | 0.345 |
| 21 | Solute carrier family 22 member 12 | 0.339 |
| 22 | Cholinesterase | 0.322 |
| 23 | Deoxynucleoside triphosphate triphosphohydrolase SAMHD1 | 0.321 |
| 24 | Microphthalmia-associated transcription factor | 0.313 |
| 25 | Ribosomal protein S6 kinase beta-1 | 0.311 |
| 26 | Solute carrier organic anion transporter family member 1B3 | 0.307 |
| 27 | Aurora kinase A | 0.302 |
| 28 | Kynurenine--oxoglutarate transaminase 3 | 0.301 |
| 29 | UDP-glucuronosyltransferase 1A6 | 0.297 |
| 30 | Palmitoleoyl-protein carboxylesterase NOTUM | 0.295 |
| 31 | Nuclear factor erythroid 2-related factor 2 | 0.295 |
| 32 | Frizzled-5 | 0.295 |
| 33 | Kynurenine--oxoglutarate transaminase 1 | 0.294 |
| 34 | Frizzled-7 | 0.293 |
| 35 | Solute carrier family 22 member 1 | 0.292 |
| 36 | UDP-glucuronosyltransferase 2B7 | 0.291 |
| 37 | Vitamin D3 receptor | 0.288 |
| 38 | Aspartate aminotransferase, cytoplasmic | 0.283 |
| 39 | UDP-glucuronosyltransferase 1A4 | 0.283 |
| 40 | UDP-glucuronosyltransferase 1A3 | 0.282 |
| 41 | Cytochrome P450 2C8 | 0.281 |
| 42 | Cytochrome P450 2B6 | 0.279 |
| 43 | Chymotrypsin-C | 0.279 |
| 44 | Solute carrier organic anion transporter family member 2B1 | 0.277 |
| 45 | UDP-glucuronosyltransferase 1A1 | 0.276 |
| 46 | Albumin | 0.272 |
| 47 | UDP-glucuronosyltransferase 1A9 | 0.270 |
| 48 | Malate dehydrogenase, cytoplasmic | 0.251 |
| 49 | ATP-binding cassette sub-family C member 2 | 0.250 |
| 50 | Serine/threonine-protein kinase PLK1 | 0.247 |
| 51 | Solute carrier family 2, facilitated glucose transporter member 1 | 0.239 |
| 52 | TAR DNA-binding protein 43 | 0.239 |
| 53 | Solute carrier family 22 member 3 | 0.234 |
| 54 | Sodium channel protein type 9 subunit alpha | 0.232 |
| 55 | Sodium channel protein type 5 subunit alpha | 0.218 |
| 56 | Sodium channel protein type 4 subunit alpha | 0.215 |
| 57 | ATPase family AAA domain-containing protein 5 | 0.210 |
| 58 | Sodium channel protein type 2 subunit alpha | 0.204 |
| 59 | ATP-dependent translocase ABCB1 | 0.204 |

Supplementary Table S2F: All 51 protein targets predicted by DRIFT for phenytoin, ranked by DRIFT score. Figure 2F shows only the highest-ranked of these.

| **Rank** | **Predicted target** | **DRIFT score** |
| --- | --- | --- |
| 1 | Sodium channel alpha subunit | 0.926 |
| 2 | Cytochrome P450 2C9 | 0.751 |
| 3 | Histamine H1 receptor | 0.741 |
| 4 | Matrix metalloproteinase-9 | 0.739 |
| 5 | 5-hydroxytryptamine receptor 2A | 0.733 |
| 6 | Sodium channel protein type 4 subunit alpha | 0.722 |
| 7 | Nuclear receptor subfamily 1 group I member 3 | 0.710 |
| 8 | Aldo-keto reductase family 1 member B1 | 0.709 |
| 9 | Sodium channel protein type 5 subunit alpha | 0.706 |
| 10 | Cannabinoid receptor 1 | 0.705 |
| 11 | Sodium channel protein type 2 subunit alpha | 0.704 |
| 12 | Cannabinoid receptor 2 | 0.703 |
| 13 | Nuclear receptor subfamily 1 group I member 2 | 0.692 |
| 14 | Cytochrome P450 2C19 | 0.690 |
| 15 | Nuclear factor erythroid 2-related factor 2 | 0.690 |
| 16 | Geminin | 0.690 |
| 17 | Solute carrier organic anion transporter family member 1B1 | 0.690 |
| 18 | Solute carrier organic anion transporter family member 1B3 | 0.690 |
| 19 | Solute carrier organic anion transporter family member 2B1 | 0.690 |
| 20 | Amine oxidase [flavin-containing] A | 0.690 |
| 21 | Fatty-acid amide hydrolase 1 | 0.690 |
| 22 | Cytochrome P450 2D6 | 0.690 |
| 23 | Albumin | 0.690 |
| 24 | Cytochrome P450 1A2 | 0.690 |
| 25 | Cytochrome P450 3A4 | 0.690 |
| 26 | Cytochrome P450 2C8 | 0.690 |
| 27 | Deoxynucleoside triphosphate triphosphohydrolase SAMHD1 | 0.690 |
| 28 | Cytochrome P450 2B6 | 0.690 |
| 29 | Solute carrier family 22 member 1 | 0.690 |
| 30 | ATP-binding cassette sub-family C member 2 | 0.690 |
| 31 | Voltage-gated inwardly rectifying potassium channel KCNH2 | 0.671 |
| 32 | ATP-dependent translocase ABCB1 | 0.671 |
| 33 | Aquaporin-4 | 0.671 |
| 34 | UDP-glucuronosyltransferase 1A1 | 0.486 |
| 35 | UDP-glucuronosyltransferase 1A4 | 0.486 |
| 36 | UDP-glucuronosyltransferase 2B7 | 0.486 |
| 37 | UDP-glucuronosyltransferase 2B15 | 0.486 |
| 38 | Integrin alpha-4 | 0.441 |
| 39 | Aldo-keto reductase family 1 member A1 | 0.380 |
| 40 | 72 kDa type IV collagenase | 0.262 |
| 41 | Multidrug and toxin extrusion protein 1 | 0.256 |
| 42 | Multidrug and toxin extrusion protein 2 | 0.256 |
| 43 | Ubiquitin carboxyl-terminal hydrolase 1 | 0.248 |
| 44 | Intermediate conductance calcium-activated potassium channel protein 4 | 0.244 |
| 45 | Multidrug resistance associated protein | 0.239 |
| 46 | Multidrug resistance-associated protein 1 | 0.239 |
| 47 | A-type voltage-gated potassium channel KCND3 | 0.229 |
| 48 | Lysine-specific demethylase 4A | 0.210 |
| 49 | Rap guanine nucleotide exchange factor 3 | 0.207 |
| 50 | Serine/threonine-protein kinase PLK1 | 0.207 |
| 51 | D-3-phosphoglycerate dehydrogenase | 0.196 |

Supplementary Table S2G: All 57 protein targets predicted by DRIFT for cenobamate, ranked by DRIFT score. Figure 2G shows only the highest-ranked of these.

| **Rank** | **Predicted target** | **DRIFT score** |
| --- | --- | --- |
| 1 | GABA-A receptor; anion channel | 0.948 |
| 2 | Sodium channel alpha subunit | 0.946 |
| 3 | Vasopressin V2 receptor | 0.486 |
| 4 | Lysophosphatidic acid receptor 1 | 0.468 |
| 5 | Vasopressin V1a receptor | 0.454 |
| 6 | Caspase-3 | 0.430 |
| 7 | Retinoic acid receptor RXR-alpha | 0.420 |
| 8 | 5-hydroxytryptamine receptor 2A | 0.409 |
| 9 | Receptor-interacting serine/threonine-protein kinase 1 | 0.369 |
| 10 | Glutaminase kidney isoform, mitochondrial | 0.357 |
| 11 | Lysophosphatidic acid receptor 3 | 0.343 |
| 12 | Acrosin | 0.327 |
| 13 | Lysophosphatidic acid receptor 2 | 0.309 |
| 14 | Serine protease 1 | 0.304 |
| 15 | Fatty-acid amide hydrolase 1 | 0.292 |
| 16 | G-protein coupled receptor 55 | 0.273 |
| 17 | Heat shock protein HSP 90-alpha | 0.260 |
| 18 | Amine oxidase [copper-containing] 3 | 0.256 |
| 19 | Monoglyceride lipase | 0.254 |
| 20 | Lysosomal protective protein | 0.252 |
| 21 | Cytochrome P450 3A4 | 0.245 |
| 22 | Cytochrome P450 2C19 | 0.236 |
| 23 | Cytochrome P450 2C9 | 0.234 |
| 24 | Glucagon-like peptide 1 receptor | 0.234 |
| 25 | Macrophage migration inhibitory factor | 0.230 |
| 26 | Histone deacetylase 2 | 0.226 |
| 27 | Receptor-interacting serine/threonine-protein kinase 3 | 0.225 |
| 28 | Tyrosyl-DNA phosphodiesterase 1 | 0.220 |
| 29 | Neutrophil elastase | 0.218 |
| 30 | Type-1 angiotensin II receptor | 0.217 |
| 31 | N-glycosylase/DNA lyase | 0.217 |
| 32 | Sphingosine 1-phosphate receptor 1 | 0.216 |
| 33 | Endonuclease 8-like 1 | 0.216 |
| 34 | Free fatty acid receptor 4 | 0.214 |
| 35 | Free fatty acid receptor 1 | 0.214 |
| 36 | Adhesion G-protein coupled receptor F1 | 0.214 |
| 37 | Endonuclease 8-like 2 | 0.213 |
| 38 | Uracil-DNA glycosylase | 0.210 |
| 39 | N-formyl peptide receptor 2 | 0.209 |
| 40 | Glucose-dependent insulinotropic receptor | 0.208 |
| 41 | C5a anaphylatoxin chemotactic receptor 1 | 0.208 |
| 42 | Beta-2 adrenergic receptor | 0.208 |
| 43 | Alpha-2A adrenergic receptor | 0.208 |
| 44 | Methyl-CpG-binding domain protein 4 | 0.207 |
| 45 | Apelin receptor | 0.207 |
| 46 | G/T mismatch-specific thymine DNA glycosylase | 0.206 |
| 47 | Serine/threonine-protein kinase PLK1 | 0.205 |
| 48 | G-protein coupled receptor 35 | 0.204 |
| 49 | CX3C chemokine receptor 1 | 0.204 |
| 50 | 5'-nucleotidase | 0.201 |
| 51 | Single-strand selective monofunctional uracil DNA glycosylase | 0.199 |
| 52 | Cytochrome P450 2D6 | 0.198 |
| 53 | DNA-3-methyladenine glycosylase | 0.198 |
| 54 | Cytochrome P450 1A2 | 0.197 |
| 55 | Voltage-gated inwardly rectifying potassium channel KCNH2 | 0.174 |
| 56 | Adenine DNA glycosylase | 0.161 |
| 57 | Guanine nucleotide-binding protein G(s) subunit alpha isoforms short | 0.153 |

Supplementary Table S3A: All 40 Reactome pathways enriched at FDR < 0.05 in the predicted target set of carbamazepine, ranked by enrichment score. FDR is the Benjamini-Hochberg adjusted p-value.

| **Rank** | **Reactome pathway** | **Pathway score** | **FDR** |
| --- | --- | --- | --- |
| 1 | Aspirin ADME | 0.900 | 9.5 × 10⁻21 |
| 2 | Interaction between L1 and Ankyrins | 0.320 | 8.1 × 10⁻15 |
| 3 | Phase 0 - rapid depolarisation | 0.320 | 8.1 × 10⁻15 |
| 4 | Glucuronidation | 0.250 | 1.0 × 10⁻8 |
| 5 | Heme degradation | 0.250 | 1.7 × 10⁻9 |
| 6 | Xenobiotics | 0.250 | 1.2 × 10⁻8 |
| 7 | Atorvastatin ADME | 0.250 | 2.7 × 10⁻11 |
| 8 | CYP2E1 reactions | 0.210 | 1.2 × 10⁻8 |
| 9 | Biosynthesis of maresin-like SPMs | 0.200 | 3.5 × 10⁻10 |
| 10 | Prednisone ADME | 0.200 | 9.1 × 10⁻9 |
| 11 | Elevation of cytosolic Ca2+ levels | 0.170 | 1.0 × 10⁻7 |
| 12 | Synthesis of epoxy (EET) and dihydroxyeicosatrienoic acids (DHET) | 0.170 | 3.5 × 10⁻7 |
| 13 | Synthesis of (16-20)-hydroxyeicosatetraenoic acids (HETE) | 0.170 | 5.8 × 10⁻7 |
| 14 | Platelet homeostasis | 0.170 | 2.3 × 10⁻9 |
| 15 | Paracetamol ADME | 0.170 | 1.0 × 10⁻4 |
| 16 | Recycling of bile acids and salts | 0.140 | 6.2 × 10⁻4 |
| 17 | Transport of organic anions | 0.140 | 1.4 × 10⁻4 |
| 18 | Neurotransmitter clearance | 0.130 | 3.8 × 10⁻6 |
| 19 | SUMOylation of intracellular receptors | 0.130 | 0.003 |
| 20 | Sensory perception of sweet, bitter, and umami (glutamate) taste | 0.130 | 0.006 |
| 21 | Fatty acids | 0.120 | 0.013 |
| 22 | Phase I - Functionalization of compounds | 0.120 | 0.021 |
| 23 | Abacavir transmembrane transport | 0.120 | 0.002 |
| 24 | Aflatoxin activation and detoxification | 0.120 | 0.019 |
| 25 | RUNX2 regulates osteoblast differentiation | 0.120 | 0.024 |
| 26 | Ciprofloxacin ADME | 0.120 | 0.002 |
| 27 | Synthesis of PC | 0.110 | 0.029 |
| 28 | Class A/1 (Rhodopsin-like receptors) | 0.110 | 0.019 |
| 29 | Synthesis, secretion, and deacylation of Ghrelin | 0.110 | 0.019 |
| 30 | Purinergic signaling in leishmaniasis infection | 0.110 | 0.028 |
| 31 | Aromatic amines can be N-hydroxylated or N-dealkylated by CYP1A2 | 0.100 | 0.019 |
| 32 | Defective UGT1A1 causes hyperbilirubinemia | 0.100 | 0.019 |
| 33 | Defective UGT1A4 causes hyperbilirubinemia | 0.100 | 0.019 |
| 34 | Defective SLCO1B3 causes hyperbilirubinemia, Rotor type (HBLRR) | 0.100 | 0.019 |
| 35 | Defective SLCO1B1 causes hyperbilirubinemia, Rotor type (HBLRR) | 0.100 | 0.019 |
| 36 | Defective ABCC2 causes DJS | 0.100 | 0.019 |
| 37 | Mitotic Metaphase/Anaphase Transition | 0.100 | 0.034 |
| 38 | Mitotic Telophase/Cytokinesis | 0.100 | 0.048 |
| 39 | Nucleotide catabolism | 0.100 | 0.019 |
| 40 | Regulation of MITF-M dependent genes involved in metabolism | 0.100 | 0.048 |

Supplementary Table S3B: All 31 Reactome pathways enriched at FDR < 0.05 in the predicted target set of lamotrigine, ranked by enrichment score. FDR is the Benjamini-Hochberg adjusted p-value.

| **Rank** | **Reactome pathway** | **Pathway score** | **FDR** |
| --- | --- | --- | --- |
| 1 | Interaction between L1 and Ankyrins | 0.900 | 4.6 × 10⁻16 |
| 2 | Phase 0 - rapid depolarisation | 0.900 | 4.6 × 10⁻16 |
| 3 | Aspirin ADME | 0.430 | 3.3 × 10⁻7 |
| 4 | Xenobiotics | 0.250 | 2.9 × 10⁻5 |
| 5 | Biosynthesis of maresin-like SPMs | 0.250 | 1.0 × 10⁻7 |
| 6 | Heme degradation | 0.240 | 7.2 × 10⁻6 |
| 7 | Sensory perception of sweet, bitter, and umami (glutamate) taste | 0.190 | 0.005 |
| 8 | Recycling of bile acids and salts | 0.180 | 4.0 × 10⁻4 |
| 9 | CYP2E1 reactions | 0.180 | 9.0 × 10⁻5 |
| 10 | Synthesis of epoxy (EET) and dihydroxyeicosatrienoic acids (DHET) | 0.180 | 4.8 × 10⁻5 |
| 11 | Synthesis of (16-20)-hydroxyeicosatetraenoic acids (HETE) | 0.180 | 5.6 × 10⁻5 |
| 12 | Transport of organic anions | 0.180 | 9.0 × 10⁻5 |
| 13 | Atorvastatin ADME | 0.180 | 5.6 × 10⁻5 |
| 14 | Voltage gated Potassium channels | 0.130 | 0.050 |
| 15 | Transport of bile salts and organic acids, metal ions and amine compounds | 0.130 | 0.006 |
| 16 | Aflatoxin activation and detoxification | 0.130 | 0.021 |
| 17 | Prednisone ADME | 0.130 | 0.006 |
| 18 | Biogenic amines are oxidatively deaminated to aldehydes by MAOA and MAOB | 0.110 | 0.037 |
| 19 | Aromatic amines can be N-hydroxylated or N-dealkylated by CYP1A2 | 0.110 | 0.021 |
| 20 | Enzymatic degradation of dopamine by COMT | 0.110 | 0.050 |
| 21 | Enzymatic degradation of Dopamine by monoamine oxidase | 0.110 | 0.037 |
| 22 | Dopamine clearance from the synaptic cleft | 0.110 | 0.037 |
| 23 | Metabolism of serotonin | 0.110 | 0.037 |
| 24 | Defective MAOA causes BRUNS | 0.110 | 0.021 |
| 25 | Defective UGT1A4 causes hyperbilirubinemia | 0.110 | 0.021 |
| 26 | Defective SLCO1B3 causes hyperbilirubinemia, Rotor type (HBLRR) | 0.110 | 0.021 |
| 27 | Defective SLCO1B1 causes hyperbilirubinemia, Rotor type (HBLRR) | 0.110 | 0.021 |
| 28 | Nucleotide catabolism | 0.110 | 0.021 |
| 29 | Defective CYP19A1 causes AEXS | 0.100 | 0.021 |
| 30 | Defective SLC2A1 causes GLUT1 deficiency syndrome 1 (GLUT1DS1) | 0.100 | 0.021 |
| 31 | Lactose synthesis | 0.100 | 0.050 |

Supplementary Table S3C: All 8 Reactome pathways enriched at FDR < 0.05 in the predicted target set of lacosamide, ranked by enrichment score. FDR is the Benjamini-Hochberg adjusted p-value.

| **Rank** | **Reactome pathway** | **Pathway score** | **FDR** |
| --- | --- | --- | --- |
| 1 | Reversible hydration of carbon dioxide | 0.900 | 1.7 × 10⁻32 |
| 2 | Phase 0 - rapid depolarisation | 0.270 | 8.4 × 10⁻22 |
| 3 | Interaction between L1 and Ankyrins | 0.210 | 1.4 × 10⁻19 |
| 4 | Erythrocytes take up carbon dioxide and release oxygen | 0.150 | 1.1 × 10⁻5 |
| 5 | Erythrocytes take up oxygen and release carbon dioxide | 0.150 | 3.5 × 10⁻6 |
| 6 | Sensory perception of sweet, bitter, and umami (glutamate) taste | 0.120 | 4.2 × 10⁻4 |
| 7 | Voltage gated Potassium channels | 0.110 | 0.015 |
| 8 | Phase 1 - inactivation of fast Na+ channels | 0.100 | 0.050 |

Supplementary Table S3D: All 40 Reactome pathways enriched at FDR < 0.05 in the predicted target set of eslicarbazepine, ranked by enrichment score. FDR is the Benjamini-Hochberg adjusted p-value.

| **Rank** | **Reactome pathway** | **Pathway score** | **FDR** |
| --- | --- | --- | --- |
| 1 | Aspirin ADME | 0.900 | 6.8 × 10⁻20 |
| 2 | Interaction between L1 and Ankyrins | 0.320 | 3.3 × 10⁻14 |
| 3 | Phase 0 - rapid depolarisation | 0.320 | 3.3 × 10⁻14 |
| 4 | Xenobiotics | 0.270 | 3.0 × 10⁻8 |
| 5 | Heme degradation | 0.230 | 4.0 × 10⁻9 |
| 6 | Atorvastatin ADME | 0.230 | 6.6 × 10⁻11 |
| 7 | Glucuronidation | 0.220 | 2.8 × 10⁻8 |
| 8 | Biosynthesis of maresin-like SPMs | 0.220 | 7.5 × 10⁻10 |
| 9 | CYP2E1 reactions | 0.210 | 3.0 × 10⁻8 |
| 10 | Prednisone ADME | 0.190 | 2.2 × 10⁻8 |
| 11 | Synthesis of epoxy (EET) and dihydroxyeicosatrienoic acids (DHET) | 0.180 | 7.7 × 10⁻7 |
| 12 | Synthesis of (16-20)-hydroxyeicosatetraenoic acids (HETE) | 0.180 | 1.3 × 10⁻6 |
| 13 | Paracetamol ADME | 0.150 | 2.0 × 10⁻4 |
| 14 | Neurotransmitter clearance | 0.140 | 7.3 × 10⁻6 |
| 15 | Phase I - Functionalization of compounds | 0.140 | 0.002 |
| 16 | SUMOylation of intracellular receptors | 0.140 | 0.005 |
| 17 | Tryptophan catabolism | 0.140 | 4.9 × 10⁻4 |
| 18 | Recycling of bile acids and salts | 0.130 | 0.001 |
| 19 | Transport of organic anions | 0.130 | 2.4 × 10⁻4 |
| 20 | Synthesis of PC | 0.120 | 0.041 |
| 21 | Fatty acids | 0.120 | 0.018 |
| 22 | Abacavir transmembrane transport | 0.120 | 1.7 × 10⁻5 |
| 23 | Synthesis, secretion, and deacylation of Ghrelin | 0.120 | 0.024 |
| 24 | Aflatoxin activation and detoxification | 0.120 | 0.024 |
| 25 | RUNX2 regulates osteoblast differentiation | 0.120 | 0.034 |
| 26 | Sensory perception of sweet, bitter, and umami (glutamate) taste | 0.120 | 0.009 |
| 27 | Organic cation transport | 0.110 | 0.009 |
| 28 | Ciprofloxacin ADME | 0.110 | 0.002 |
| 29 | Malate-aspartate shuttle | 0.110 | 0.007 |
| 30 | Aromatic amines can be N-hydroxylated or N-dealkylated by CYP1A2 | 0.100 | 0.024 |
| 31 | Defective UGT1A1 causes hyperbilirubinemia | 0.100 | 0.024 |
| 32 | Defective UGT1A4 causes hyperbilirubinemia | 0.100 | 0.024 |
| 33 | Defective SLC2A1 causes GLUT1 deficiency syndrome 1 (GLUT1DS1) | 0.100 | 0.024 |
| 34 | Defective SLCO1B3 causes hyperbilirubinemia, Rotor type (HBLRR) | 0.100 | 0.024 |
| 35 | Defective SLC22A12 causes renal hypouricemia 1 (RHUC1) | 0.100 | 0.024 |
| 36 | Defective SLCO1B1 causes hyperbilirubinemia, Rotor type (HBLRR) | 0.100 | 0.024 |
| 37 | Defective ABCC2 causes DJS | 0.100 | 0.024 |
| 38 | Mitotic Metaphase/Anaphase Transition | 0.100 | 0.042 |
| 39 | Interaction between PHLDA1 and AURKA | 0.100 | 0.042 |
| 40 | Nucleotide catabolism | 0.100 | 0.024 |

Supplementary Table S3E: All 40 Reactome pathways enriched at FDR < 0.05 in the predicted target set of licarbazepine, ranked by enrichment score. FDR is the Benjamini-Hochberg adjusted p-value.

| **Rank** | **Reactome pathway** | **Pathway score** | **FDR** |
| --- | --- | --- | --- |
| 1 | Aspirin ADME | 0.900 | 6.8 × 10⁻20 |
| 2 | Interaction between L1 and Ankyrins | 0.320 | 3.3 × 10⁻14 |
| 3 | Phase 0 - rapid depolarisation | 0.320 | 3.3 × 10⁻14 |
| 4 | Xenobiotics | 0.270 | 3.0 × 10⁻8 |
| 5 | Heme degradation | 0.230 | 4.0 × 10⁻9 |
| 6 | Atorvastatin ADME | 0.230 | 6.6 × 10⁻11 |
| 7 | Glucuronidation | 0.220 | 2.8 × 10⁻8 |
| 8 | Biosynthesis of maresin-like SPMs | 0.220 | 7.5 × 10⁻10 |
| 9 | CYP2E1 reactions | 0.210 | 3.0 × 10⁻8 |
| 10 | Prednisone ADME | 0.190 | 2.2 × 10⁻8 |
| 11 | Synthesis of epoxy (EET) and dihydroxyeicosatrienoic acids (DHET) | 0.180 | 7.7 × 10⁻7 |
| 12 | Synthesis of (16-20)-hydroxyeicosatetraenoic acids (HETE) | 0.180 | 1.3 × 10⁻6 |
| 13 | Paracetamol ADME | 0.150 | 2.0 × 10⁻4 |
| 14 | Neurotransmitter clearance | 0.140 | 7.3 × 10⁻6 |
| 15 | Phase I - Functionalization of compounds | 0.140 | 0.002 |
| 16 | SUMOylation of intracellular receptors | 0.140 | 0.005 |
| 17 | Tryptophan catabolism | 0.140 | 4.9 × 10⁻4 |
| 18 | Recycling of bile acids and salts | 0.130 | 0.001 |
| 19 | Transport of organic anions | 0.130 | 2.4 × 10⁻4 |
| 20 | Synthesis of PC | 0.120 | 0.041 |
| 21 | Fatty acids | 0.120 | 0.018 |
| 22 | Abacavir transmembrane transport | 0.120 | 1.7 × 10⁻5 |
| 23 | Synthesis, secretion, and deacylation of Ghrelin | 0.120 | 0.024 |
| 24 | Aflatoxin activation and detoxification | 0.120 | 0.024 |
| 25 | RUNX2 regulates osteoblast differentiation | 0.120 | 0.034 |
| 26 | Sensory perception of sweet, bitter, and umami (glutamate) taste | 0.120 | 0.009 |
| 27 | Organic cation transport | 0.110 | 0.009 |
| 28 | Ciprofloxacin ADME | 0.110 | 0.002 |
| 29 | Malate-aspartate shuttle | 0.110 | 0.007 |
| 30 | Aromatic amines can be N-hydroxylated or N-dealkylated by CYP1A2 | 0.100 | 0.024 |
| 31 | Defective UGT1A1 causes hyperbilirubinemia | 0.100 | 0.024 |
| 32 | Defective UGT1A4 causes hyperbilirubinemia | 0.100 | 0.024 |
| 33 | Defective SLC2A1 causes GLUT1 deficiency syndrome 1 (GLUT1DS1) | 0.100 | 0.024 |
| 34 | Defective SLCO1B3 causes hyperbilirubinemia, Rotor type (HBLRR) | 0.100 | 0.024 |
| 35 | Defective SLC22A12 causes renal hypouricemia 1 (RHUC1) | 0.100 | 0.024 |
| 36 | Defective SLCO1B1 causes hyperbilirubinemia, Rotor type (HBLRR) | 0.100 | 0.024 |
| 37 | Defective ABCC2 causes DJS | 0.100 | 0.024 |
| 38 | Mitotic Metaphase/Anaphase Transition | 0.100 | 0.042 |
| 39 | Interaction between PHLDA1 and AURKA | 0.100 | 0.042 |
| 40 | Nucleotide catabolism | 0.100 | 0.024 |

Supplementary Table S3F: All 45 Reactome pathways enriched at FDR < 0.05 in the predicted target set of phenytoin, ranked by enrichment score. FDR is the Benjamini-Hochberg adjusted p-value.

| **Rank** | **Reactome pathway** | **Pathway score** | **FDR** |
| --- | --- | --- | --- |
| 1 | Aspirin ADME | 0.900 | 8.3 × 10⁻15 |
| 2 | Interaction between L1 and Ankyrins | 0.420 | 8.3 × 10⁻15 |
| 3 | Phase 0 - rapid depolarisation | 0.420 | 8.3 × 10⁻15 |
| 4 | Heme degradation | 0.390 | 1.6 × 10⁻11 |
| 5 | Xenobiotics | 0.370 | 1.7 × 10⁻8 |
| 6 | CYP2E1 reactions | 0.290 | 1.7 × 10⁻8 |
| 7 | Biosynthesis of maresin-like SPMs | 0.290 | 3.6 × 10⁻10 |
| 8 | Atorvastatin ADME | 0.280 | 6.2 × 10⁻9 |
| 9 | Synthesis of epoxy (EET) and dihydroxyeicosatrienoic acids (DHET) | 0.220 | 5.1 × 10⁻7 |
| 10 | Synthesis of (16-20)-hydroxyeicosatetraenoic acids (HETE) | 0.220 | 8.3 × 10⁻7 |
| 11 | Prednisone ADME | 0.210 | 1.3 × 10⁻6 |
| 12 | Glucuronidation | 0.180 | 3.8 × 10⁻5 |
| 13 | Paracetamol ADME | 0.180 | 1.3 × 10⁻4 |
| 14 | Recycling of bile acids and salts | 0.160 | 7.6 × 10⁻4 |
| 15 | Transport of organic anions | 0.160 | 1.7 × 10⁻4 |
| 16 | ABC-family proteins mediated transport | 0.150 | 0.028 |
| 17 | Interleukin-4 and Interleukin-13 signaling | 0.150 | 0.050 |
| 18 | Norepinephrine Neurotransmitter Release Cycle | 0.130 | 0.021 |
| 19 | Fatty acids | 0.130 | 0.015 |
| 20 | Phase I - Functionalization of compounds | 0.130 | 0.022 |
| 21 | Abacavir transmembrane transport | 0.130 | 0.002 |
| 22 | Class A/1 (Rhodopsin-like receptors) | 0.130 | 0.021 |
| 23 | Aflatoxin activation and detoxification | 0.130 | 0.021 |
| 24 | Sensory perception of sweet, bitter, and umami (glutamate) taste | 0.130 | 0.007 |
| 25 | Ciprofloxacin ADME | 0.130 | 0.002 |
| 26 | Activation of Matrix Metalloproteinases | 0.120 | 0.037 |
| 27 | NFE2L2 regulating MDR associated enzymes | 0.120 | 0.005 |
| 28 | Biogenic amines are oxidatively deaminated to aldehydes by MAOA and MAOB | 0.110 | 0.033 |
| 29 | Aromatic amines can be N-hydroxylated or N-dealkylated by CYP1A2 | 0.110 | 0.021 |
| 30 | Enzymatic degradation of dopamine by COMT | 0.110 | 0.045 |
| 31 | Enzymatic degradation of Dopamine by monoamine oxidase | 0.110 | 0.033 |
| 32 | Dopamine clearance from the synaptic cleft | 0.110 | 0.033 |
| 33 | Metabolism of serotonin | 0.110 | 0.033 |
| 34 | Transport of bile salts and organic acids, metal ions and amine compounds | 0.110 | 0.008 |
| 35 | Defective MAOA causes BRUNS | 0.110 | 0.021 |
| 36 | Defective SLCO1B3 causes hyperbilirubinemia, Rotor type (HBLRR) | 0.110 | 0.021 |
| 37 | Defective SLCO1B1 causes hyperbilirubinemia, Rotor type (HBLRR) | 0.110 | 0.021 |
| 38 | Fructose biosynthesis | 0.110 | 0.033 |
| 39 | Defective ABCC2 causes DJS | 0.110 | 0.021 |
| 40 | Nucleotide catabolism | 0.110 | 0.021 |
| 41 | NFE2L2 regulating TCA cycle genes | 0.110 | 0.045 |
| 42 | Defective UGT1A1 causes hyperbilirubinemia | 0.100 | 0.021 |
| 43 | Defective UGT1A4 causes hyperbilirubinemia | 0.100 | 0.021 |
| 44 | Mitotic Metaphase/Anaphase Transition | 0.100 | 0.033 |
| 45 | Mitotic Telophase/Cytokinesis | 0.100 | 0.045 |

Supplementary Table S3G: All 28 Reactome pathways enriched at FDR < 0.05 in the predicted target set of cenobamate, ranked by enrichment score. FDR is the Benjamini-Hochberg adjusted p-value.

| **Rank** | **Reactome pathway** | **Pathway score** | **FDR** |
| --- | --- | --- | --- |
| 1 | G alpha (i) signalling events | 0.900 | 2.2 × 10⁻4 |
| 2 | G alpha (q) signalling events | 0.890 | 7.7 × 10⁻5 |
| 3 | Interaction between L1 and Ankyrins | 0.390 | 6.2 × 10⁻13 |
| 4 | Phase 0 - rapid depolarisation | 0.390 | 6.2 × 10⁻13 |
| 5 | GABA receptor activation | 0.360 | 7.1 × 10⁻21 |
| 6 | Recognition and association of DNA glycosylase with site containing an affected pyrimidine | 0.330 | 1.0 × 10⁻5 |
| 7 | Cleavage of the damaged pyrimidine | 0.330 | 1.0 × 10⁻5 |
| 8 | Displacement of DNA glycosylase by APEX1 | 0.260 | 1.6 × 10⁻7 |
| 9 | Lysosphingolipid and LPA receptors | 0.260 | 5.4 × 10⁻5 |
| 10 | Xenobiotics | 0.210 | 3.0 × 10⁻4 |
| 11 | Aspirin ADME | 0.210 | 0.003 |
| 12 | Biosynthesis of maresin-like SPMs | 0.200 | 1.3 × 10⁻6 |
| 13 | Signaling by ERBB4 | 0.190 | 5.4 × 10⁻9 |
| 14 | Regulation of necroptotic cell death | 0.180 | 0.008 |
| 15 | CYP2E1 reactions | 0.160 | 7.6 × 10⁻4 |
| 16 | Synthesis of epoxy (EET) and dihydroxyeicosatrienoic acids (DHET) | 0.160 | 3.0 × 10⁻4 |
| 17 | Synthesis of (16-20)-hydroxyeicosatetraenoic acids (HETE) | 0.160 | 4.2 × 10⁻4 |
| 18 | APEX1-Independent Resolution of AP Sites via the Single Nucleotide Replacement Pathway | 0.160 | 2.2 × 10⁻4 |
| 19 | Synthesis, secretion, and inactivation of Glucagon-like Peptide-1 (GLP-1) | 0.150 | 0.005 |
| 20 | Vasopressin-like receptors | 0.150 | 0.008 |
| 21 | TRIF-mediated programmed cell death | 0.130 | 0.016 |
| 22 | TLR3-mediated TICAM1-dependent programmed cell death | 0.130 | 0.008 |
| 23 | Microbial modulation of RIPK1-mediated regulated necrosis | 0.130 | 0.019 |
| 24 | SARS-CoV-1-mediated effects on programmed cell death | 0.130 | 0.008 |
| 25 | Adrenoceptors | 0.120 | 0.016 |
| 26 | Synthesis, secretion, and inactivation of Glucose-dependent Insulinotropic Polypeptide (GIP) | 0.120 | 0.030 |
| 27 | Free fatty acid receptors | 0.120 | 0.006 |
| 28 | Sensory perception of sweet, bitter, and umami (glutamate) taste | 0.120 | 0.025 |
